# Cytoneme feedback ensures signaling specificity when multiple ligands converge on a common receptor

**DOI:** 10.1101/2025.08.28.672500

**Authors:** Akshay Patel, Molly Maranto, Hind Hitti, Sougata Roy

**Affiliations:** Department of Cell Biology and Molecular Genetics, University of Maryland, College Park, Maryland, USA; National Center for Advancing Translational Sciences (NCATS), Stem Cell Translation Laboratory (SCTL), National Institutes of Health (NIH), Rockville, MD 20850, USA; Department of Pharmacology, Physiology, and Neuroscience, University of South Carolina School of Medicine, Columbia, SC USA 29209

**Keywords:** Cytonemes, polarity, positive feedback, Pointed, FGF, Heartless, Pyramus, Thisbe, stem cell niche, adult muscle precursor

## Abstract

Cell-cell communication via conserved signaling pathways often involves multiple paracrine ligands that act through shared receptors and/or downstream cascades yet elicit distinct outcomes, raising the question of how specificity is achieved. In the *Drosophila* adult-muscle-precursor (AMP) niche, the FGF ligands Pyramus and Thisbe both signal through the Heartless (Htl) receptor in isogenic AMPs yet drive divergent responses. *In-vivo* imaging of endogenous Pyramus:mCherry^endo^ and Thisbe:sfGFP^endo^ knock-ins revealed that the ligands segregate into distinct, receptor-bound territories among isogenic AMPs. This segregation arises as AMPs acquire ligands via polarized, Htl-containing cytonemes that directly contact each ligand source. Cytoneme polarity and target-specificity are position-dependent, enabling ligand distribution to scale with tissue size and AMP organization. A positive feedback loop through the Htl-Pointed axis reinforces cytoneme polarity, amplifying ligand-specific segregation and signaling. These findings establish cell-intrinsic feedback as a general principle for sustaining cytoneme polarity and specificity, and reveal how cytonemes confer signaling specificity to pathways activated by multiple ligands within the same progenitor population.

## INTRODUCTION

During development and tissue homeostasis, secreted developmental signals generate diverse cell types and architectures, despite often acting through a limited set of conserved pathways (Perrimon et al., 2012; Housden and Perrimon, 2014). Many growth factors, including epidermal growth factors (EGF) and fibroblast growth factors (FGF), converge on the Mitogen-activated protein kinase (MAPK) pathway (Perrimon et al., 2012). In vertebrates, 22 FGFs act through only four receptors, all signaling via MAPK (Beenken and Mohammadi, 2009; Dorey and Amaya, 2010; Itoh and Ornitz, 2004). Likewise, in *Drosophila*, the FGF8 homologs Pyramus (Pyr) and Thisbe (Ths) both signal through a common FGF receptor, Heartless (Htl), but nonetheless direct non-overlapping functions during mesodermal development (Stepanik et al., 2020; McMahon et al., 2010; Clark et al., 2011; Patel et al., 2022). Such situations illustrate the challenge of maintaining distinct, ligand-specific codes, particularly when multiple ligands disperse through a shared extracellular environment and converge on a common receptor or signaling pathway within the same progenitor populations.

One mechanism that can confer ligand specificity is cytoneme-mediated, polarized delivery of signals directly to individual target cells. Cytonemes are thin (∼100-150 nm diameter) actin-rich polarized filopodia that enable direct, target-specific presentation and exchange of developmental signals via synapse-like contacts (Kornberg, 2016; Kornberg and Roy, 2014a; Roy et al., 2014; Fereres et al., 2019; Kornberg and Roy, 2014b; Patel et al., 2022; Wood et al., 2021; Du et al., 2022). Cytonemes mediate the exchange of key ligands, including Hedgehog (Hh), Decapentaplegic (Dpp), Fibroblast Growth Factor (FGF), Epidermal Growth Factor (EGF), Ephrin, Wnt, and Notch/Delta, in both vertebrate and invertebrate systems (Zhang and Scholpp, 2019; Bischoff et al., 2013; Kornberg and Roy, 2014c; Du et al., 2018a; Chen et al., 2017; Hall et al., 2021; Huang and Kornberg, 2015; Roy et al., 2011a; Stanganello et al., 2015; Sanders et al., 2013; Krupke et al., 2016; Roy et al., 2014; Clements et al., 2023; Zhang et al., 2024; Belian et al., 2023; Wang et al., 2025; Greicius et al., 2025; Lalioti et al., 2025; Hall et al., 2024; Clements et al., 2024; Sutcliffe et al., 2025). These studies highlight cytonemes as a widely used mechanism for ensuring signaling precision and specificity in development and stem cell maintenance. Mechanistically, unlike diffusion-based ligand spread, cytoneme-mediated signaling is receptor-dependent, contact-driven (Kornberg, 2016, 2014). Cells can sort different receptors to different ligand-specific cytonemes to receive different signals (Roy et al., 2011b). However, the receptor-dependency of ligand transport together with inherent plasticity of cytonemes, invites a critical question: how do cytonemes preserve or adjust their signal- and target-specificity and navigational polarity within a noisy signaling microenvironment, particularly when multiple ligands share the same receptor and signaling pathway?

Work in the *Drosophila* air sac primordium (ASP), which receives an FGF, Branchless (Bnl) via cytonemes, demonstrated that Bnl/FGF signaling can feedback regulate Bnl-receiving cytoneme behavior, thereby sustaining target-specificity and robustness of Bnl/FGF dispersion patterns (Du et al., 2018a). Feedback loops are integral features of all signaling pathways, implicated in the spatial and temporal refinement of signaling outputs, including cell polarity and shape (Neben et al., 2019; Perrimon and McMahon, 1999; Freeman, 2000). These precedents suggest that signaling feedback-driven autoregulation of polarity may represent a core mechanistic module of cytonemes for sustaining selective targeting of distinct signaling partners and enabling context-dependent ligand dispersion. However, the applicability of this model might be limited to a single ligand-receptor pair, as observed in the ASP. It remains unknown whether such autoregulation can maintain cytoneme specificity in more complex microenvironments where multiple ligands converge on the same receptor and signaling pathway to produce distinct outcomes.

The *Drosophila* adult muscle precursor (AMP) niche in the larval wing imaginal disc offers an ideal system to address this problem. The wing disc epithelium expresses Pyr and Ths, both of which signal through Htl in AMPs, but promote distinct AMP fate choices and organization (Stathopoulos et al., 2004; Stepanik et al., 2020; Patel et al., 2022; Kadam et al., 2009b). AMPs selectively occupy FGF-expressing niches by extending Htl-containing cytonemes that extend toward the apical junctions of FGF-producing epithelium, anchoring to narrow intercellular spaces (Patel et al., 2022). FGF overexpression experiments suggested that niche-adhering AMP cytonemes can directly receive Pyr and Ths in a polarized manner, segregating them into distinct AMPs territories, even under excess ligand supply (Patel et al., 2022). Moreover, AMPs that received Pyr adopted the direct flight muscle (DFM) lineage, whereas those that received Ths adopted the indirect flight muscle (IFM) lineage. Thus, the AMP niche exemplifies how the same FGF-receptor can direct ligand-specific lineage outcomes.

The AMP niche also exhibits structural asymmetry, recapitulating critical events of stem cell niches. AMPs originate from common embryonic progenitors and attach to the FGF-expressing niche to transiently expand their population (Gunage et al., 2017). Niche-adhered AMPs divide asymmetrically, orienting their division axis orthogonal to the wing disc plane so that one daughter exits the niche and prepares for differentiation, while the other remains niche-adhered and mitotically active (Gunage et al., 2017). This division pattern generates a multi-layered AMP arrangement perpendicular to the wing disc, with FGF-responsive mitotic AMPs closest to the niche and their non-mitotic, non-FGF-signaling descendants positioned in more distal layers. Such cellular and signaling asymmetries make the AMP niche an ideal system for investigating how cytoneme-mediated signaling preserves polarity, specificity, and precision for multiple ligands by engaging the same receptor.

Here, using fully functional, genome-edited *pyr:mCherry*^endo^ and *ths:sfGFP^endo^* knock-in constructs combined with high-resolution quantitative in vivo imaging, we mapped the endogenous, non-overlapping distribution of Pyr and Ths, showing that cytonemes are essential for their segregation into distinct niche- and recipient-specific territories. Importantly, we discovered that the AMP cytoneme polarity and specificity are position-dependent, tunable functions that can scale FGF dispersion to the tissue size. We uncovered an AMP-intrinsic positive feedback loop, downstream of Htl signaling, that reinforces ligand-specific signaling precision and polarity of cytonemes and amplifies the localized signaling polarity into large-scale AMP-specific segregated Pyr and Ths signaling patterns. These findings suggest cell-intrinsic signaling feedback as a generalizable principle by which cytonemes autoregulate the specificity and precision of their contact-dependent communication for multiple ligands, when engaging the same receptor in the same progenitor population.

## RESULTS

### Defining functional tagging sites for Pyr and Ths

To visualize the native distribution patterns of Pyr and Ths, we used a GFP- or mCherry-based protein tagging strategy, expressing the tagged proteins at physiological levels from genome-edited *pyr* and *ths* knock-in alleles. As a first step, we identified internal sites in each protein backbone that were permissive for insertion of the ∼27 kDa super-folder GFP (sfGFP) or mCherry. We generated transgenic flies harboring *UAS*-FGF:sfGFP chimeras with GFP fused at different internal positions and assessed their signaling activity by ectopically expressing them under *dpp-Gal4* control. In wild-type discs, AMPs occupy only the FGF-producing notum and hinge regions of the wing imaginal disc, but not the pouch (Fig.1A,A’,C) (Patel et al., 2022; Fernandes et al., 1991; Gunage et al., 2017, 2014). However, ectopic expression of Pyr or Ths in the AMP-free wing disc pouch under *dpp-Gal4* induces AMPs to colonize the ectopic FGF-producing site via cytonemes and adopt distinct, DFM and IFM specific lineages in response to receiving Pyr and Ths, respectively (Patel et al., 2022). We took advantage of this assay, creating the ectopic FGF-producing AMP niche in the wing disc pouch, to test whether the chimeric FGF:sfGFPs could induce AMP colonization, FGF reception, and thereby demonstrate their functionality.

**Figure 1.**
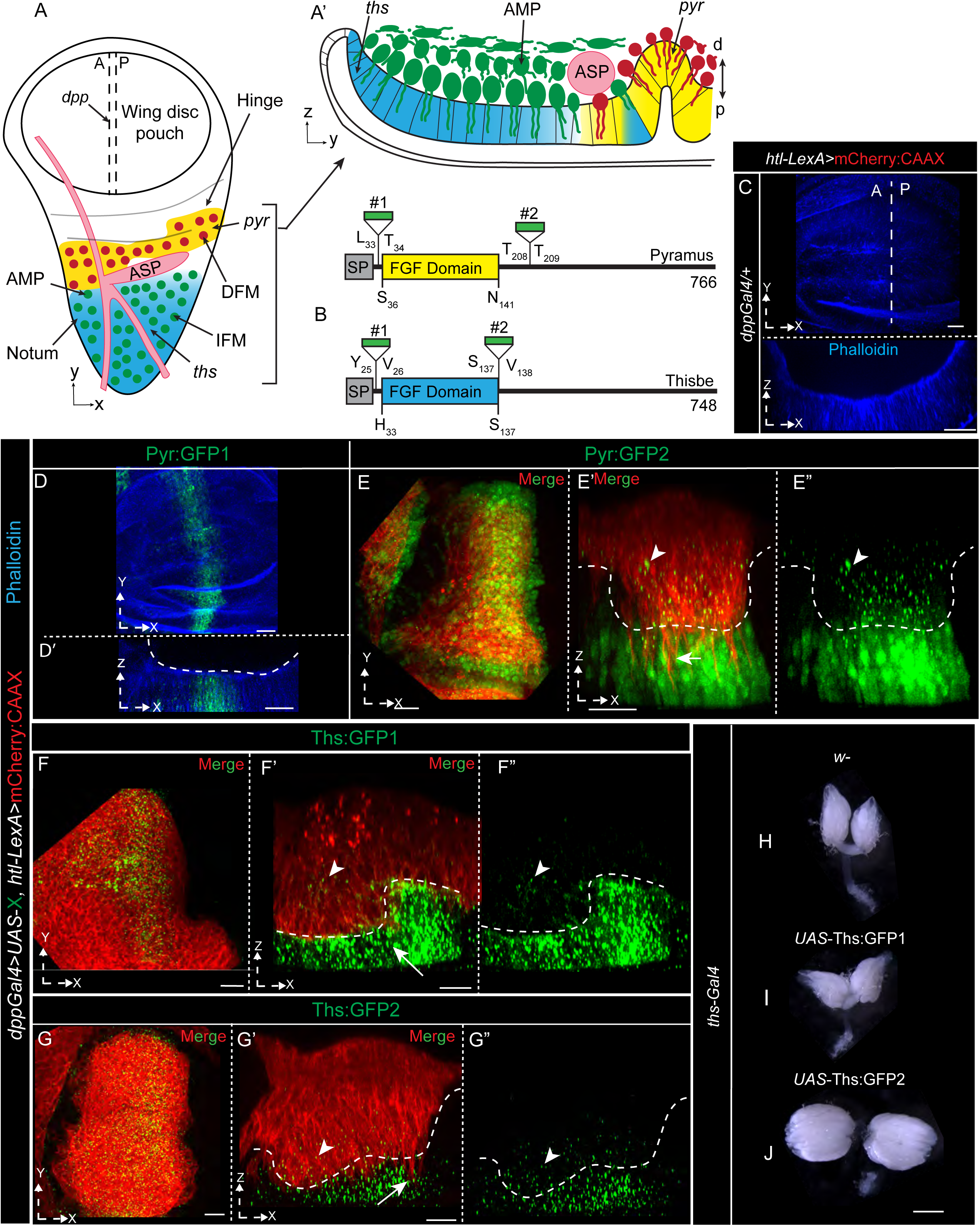
Defining functional tagging sites for Pyr and Ths. **(A-A”’)** Schematic of an L3 wing disc showing AMP organization within Pyr-(yellow) and Ths-(blue)-expressing wing disc niches in XY (A) and YZ (A’) orientations. Red AMPs: DFM lineage; green AMPs: IFM lineage; ASP: disc-associated air sac primordium. Double dashed lines in (A) mark the approximate *dpp*-expression zone at the anterior/posterior (A/P) boundary in the wing disc pouch, used as an ectopic FGF-producing niche in panels C-G”. **(B)** Domain organization of Pyr and Ths showing SP (signal peptide), conserved FGF domain, and tagging sites 1 and 2; numbers indicate amino acid positions (see Supplementary Figure 1G,H). **(C)** Phalloidin-stained (blue) wild-type wing disc pouch in XY (top) and XZ (bottom) orientations showing absence of AMPs (see Supplementary Fig. 1A-D’). **(D-G’)** Representative pouch regions expressing indicated FGF:GFP chimeras under *dpp*-*Gal4*. AMPs are marked by mCherryCAAX (red). Arrowheads: FGF:GFP puncta received by AMPs; arrows: cytonemes; dashed line: AMP-niche junction. Disc orientations, merged/split channels, and genotypes are indicated in each panel. **(H-J)** Ovary size in adult females of indicated genotypes (see Supplementary Fig. 1E,F). Also see ‘source data’ file-FigS2D for the null rescue analyses. Scale bars: 10 μm (C-H’); 0.5 mm (H-J).

Pyr is a 766-amino-acid protein that includes an N-terminal 29-residue signal peptide (SP) and a conserved FGF domain spanning amino acids 36 to 141 (Fig. 1B)(Tulin and Stathopoulos, 2010; Stathopoulos et al., 2004). We compared two sfGFP-tagged Pyr constructs (see Methods, Fig. 1B, Supplementary Fig. 1A): (i) *UAS*-Pyr:GFP1, with sfGFP inserted between L_33_ and T_34_, downstream of the SP and upstream of the FGF domain; and (ii) *UAS*-Pyr:GFP2, previously reported(Patel et al., 2022) to have sfGFP inserted between T_208_ and T_209_, downstream of the FGF domain. Similarly, Ths is a 748-amino-acid protein with an N-terminal 20-residue SP and an FGF domain spanning amino acids 33 to 137 (Tulin and Stathopoulos, 2010; Stathopoulos et al., 2004). We compared two sfGFP-tagged Ths constructs (see Methods, Figure 1B, Supplementary Fig. 1B): (i) *UAS*-Ths:GFP1, with sfGFP inserted between Y_26_ and V_27_, downstream of the SP and upstream of the FGF domain; and (ii) *UAS*-Ths:GFP2, previously reported (Patel et al., 2022) to have sfGFP inserted between S_137_ and V_138_, downstream of the FGF domain.

The expression of both Ths:GFP1 and Ths:GFP2 by *dpp-Gal4* induced AMP colonization in the ectopic Ths-expressing niche and activated MAPK signaling in niche-adhered AMPs. In contrast, only Pyr:GFP2, but not Pyr:GFP1, elicited this expected FGF gain-of-function (GOF) phenotype (Figs. 1D-G; S1A-B’). Consistent with these results, overexpression of either Ths:GFP1 or Ths:GFP2 under *ths-Gal4* (a *ths* null allele) or Pyr:GFP2 under *pyr-Gal4* (a *pyr* null allele) in flies also harboring the double *pyr* and *ths* deficient *Df(2R)BSC25* allele, rescued the lethality of *Df*(2R)BSC25/*pyr/ths*-*Gal4* trans-heterozygotes. In contrast, overexpression of Pyr:GFP1 failed to rescue the lethality of *Df*(2R)BSC25/*pyr-Gal4* animals, indicating that only tagging site 2 in Pyr is suitable for GFP insertion. Moreover, as shown in Supplementary Fig. 1C–D′, we generated a Pyr:mCherry2 construct by replacing sfGFP with mCherry at site 2. The ectopic Pyr:mCherry2-expressing niche supported AMP colonization via cytoneme-mediated ligand uptake, similar to Pyr:sfGFP2 (Fig.S1C-D’ and (Patel et al., 2022)). These results demonstrate that the functional integrity of chimeric Pyr is not influenced by the fluorescent protein.

Surprisingly, although previous results suggested that both Ths:GFP1 and Ths:GFP2 are functional ligands in the wing disc, adult females were sterile when overexpressing Ths:GFP2, but not Ths:GFP1, under *ths-Gal4* (Fig. S1E,F). These sterile females had markedly enlarged ovaries compared to wild-type (*w*^⁻^) females or females expressing Ths:GFP1 at comparable levels (Figs. 1H-J; S1E,F). These FGF gain-of-function defects are consistent with the roles of Ths in ovarian development and the maintenance of female fertility(Irizarry and Stathopoulos, 2015). These results suggest that Ths:GFP1 may function as a relatively weaker ligand than Ths:GFP2 in adult ovarian follicles. Structural predictions for both Pyr and Ths (Fig. S1G,H) indicate that tagging site 2 is located within an unstructured region, C-terminal to the conserved ‘FGF domain’ of both FGFs. Inserting a large sfGFP tag at site 2 may produce a more optimal conformation for FGFs function.

### Non-overlapping patterns of native Pyr and Ths dispersion

Based on the above results, we selected site-2 within both Pyr and Ths to generate *pyr:mCherry^endo^* and *ths:sfGFP^endo^* knock-in lines using CRISPR/Cas9-mediated genome editing method (Figs.2A,B; Supplementary Fig. 2A,B,D; see Materials and Methods). Sequence verification confirmed the successful generation of two *pyr:mCherry^endo^* and three *ths:sfGFP^endo^*lines, all of which are homozygous viable. Furthermore, trans-heterozygous animals, *pyr:mCherry^endo^*/ *pyr* null (*Df(2R)BSC25* or *pyr-Gal4*) and *ths:sfGFP^endo^* / *ths* null (*Df(2R)BSC25* or *Ths^759^* or *ths-Gal4*) were viable and displayed no morphological defects, indicating that the knock-in alleles produce fully functional proteins.

Despite imaging with a high-resolution confocal microscope equipped with a sensitive Airyscan detector, Ths:sfGFP^endo^ and Pyr:mCherry^endo^ were barely visible in the ligand-producing wing disc cells. Instead, large amounts of tagged FGFs appeared as fluorescent puncta within disc-adhered AMPs, indicating cell-non-autonomous dispersion of these ligands (Fig. 2C-J). As shown in these figures (Fig.2C-J), the non-cell-autonomous distribution of Ths:sfGFP^endo^ and Pyr:mCherry^endo^ from the wing disc to recipient AMPs was confirmed by selectively labeling either ligand-producing wing disc cells or AMPs or both in the knock-in animals. This primarily recipient-specific steady-state distribution of both ligands closely resembled previously reported native FGF/Bnl dispersion (Du et al., 2018a), suggesting that these potent growth factors may be expressed at very low levels and rapidly get transferred to recipient cells upon externalization. These analyses also revealed several notable features in Pyr and Ths dispersion.

**Figure 2.**
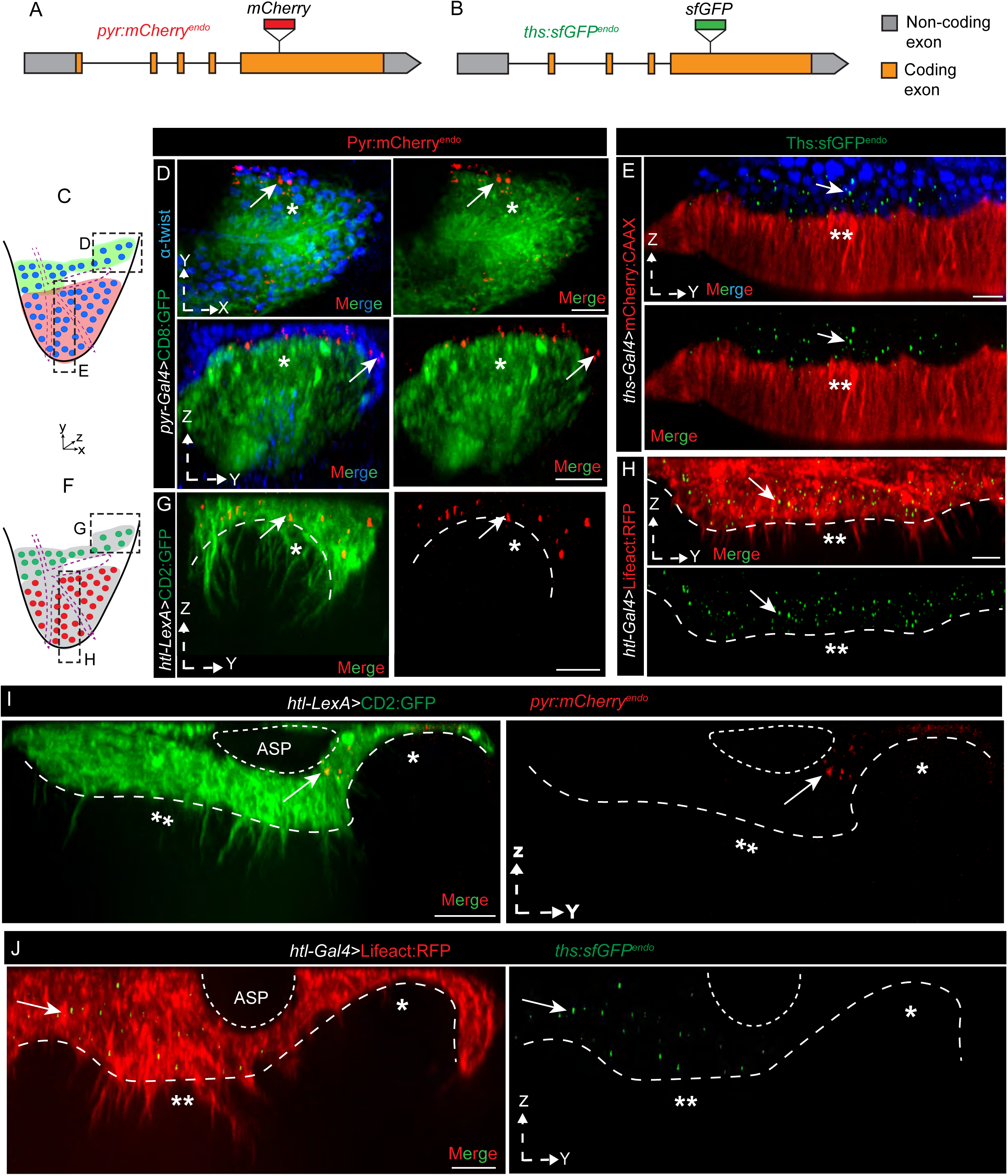
Distinct AMP-specific distribution patterns of endogenous Pyr and Ths. **(A,B)** Schematic maps (not to scale) of the *pyr*:*mCherry^endo^*and *ths*:*sfGFP^endo^* genomic loci showing in-frame tag insertion sites corresponding to site-2 in Figure 1B (see also Supplementary Fig. 2A,B,D). **(C-J)** Confocal XY or YZ views showing niche-specific (C-E,I,J), and/or AMP-specific (F-J), non-cell-autonomous dispersion of Pyr:mCherry^endo^ and Ths:sfGFP^endo^ (as indicated) from *pyr*- or *ths*-expressing wing disc sources to AMPs. **(C-E)** wing disc *pyr* and *ths* sources marked by *pyr*-*Gal4* or *ths*-*Gal4*-driven fluorescent proteins expression and AMP nuclei marked by Anti-Twist staining. **(F-J)** AMPs marked by *htl-Gal4-*driven fluorescent proteins as indicated. **(C,F)** Schematic illustrating results for D,E (C) and G,H (F-DFM precursors (green) and IFM precursors (red)); Arrows: native FGF puncta within AMP layers. *: *pyr*-expressing niche; **: *ths*-expressing niche. XY/YZ orientations, genotypes, and color codes are indicated in each panel. Dashed lines: AMP-wing disc boundary; dashed boxes in C,F: ROIs for indicated image panels. Scale bars: 10µm.

First, as shown in Figures 2I and 2J, despite being secreted, the distribution of both ligands was spatially restricted to the niche-adhering AMPs. AMPs in non-overlapping *pyr*- or *ths*-expressing wing disc niches received exclusively either Pyr or Ths, depending on which ligand-expressing domain they occupied. This specificity and segregation were maintained despite the close proximity of AMPs, their same clonal origin, and similar expression levels of Htl (Fig. 2D-G’). Second, extracellular pools of these secreted ligands detected using non-permeabilized α-GFP and α-mCherry immunostaining (α-GFP^NP^ and α-mCherry^NP^; see Materials and Methods) confirmed strictly niche-limited and non-overlapping dispersion of the chimeric FGFs along the AMP surfaces (Fig. 3A-C). Notably, some areas of wing discs, such as those beneath the air sac primordium (ASP; Fig. 1A’), contain overlapping *ths* and *pyr* expressing cells (Patel et al., 2022). AMPs in these overlapping signaling zones received both externalized Pyr:mCherry^endo^ and Ths:sfGFP^endo^ (Fig. 3A-C), indicating that AMPs can receive both ligands, but ligand reception is determined by their adherence to a specific niche.

**Figure 3.**
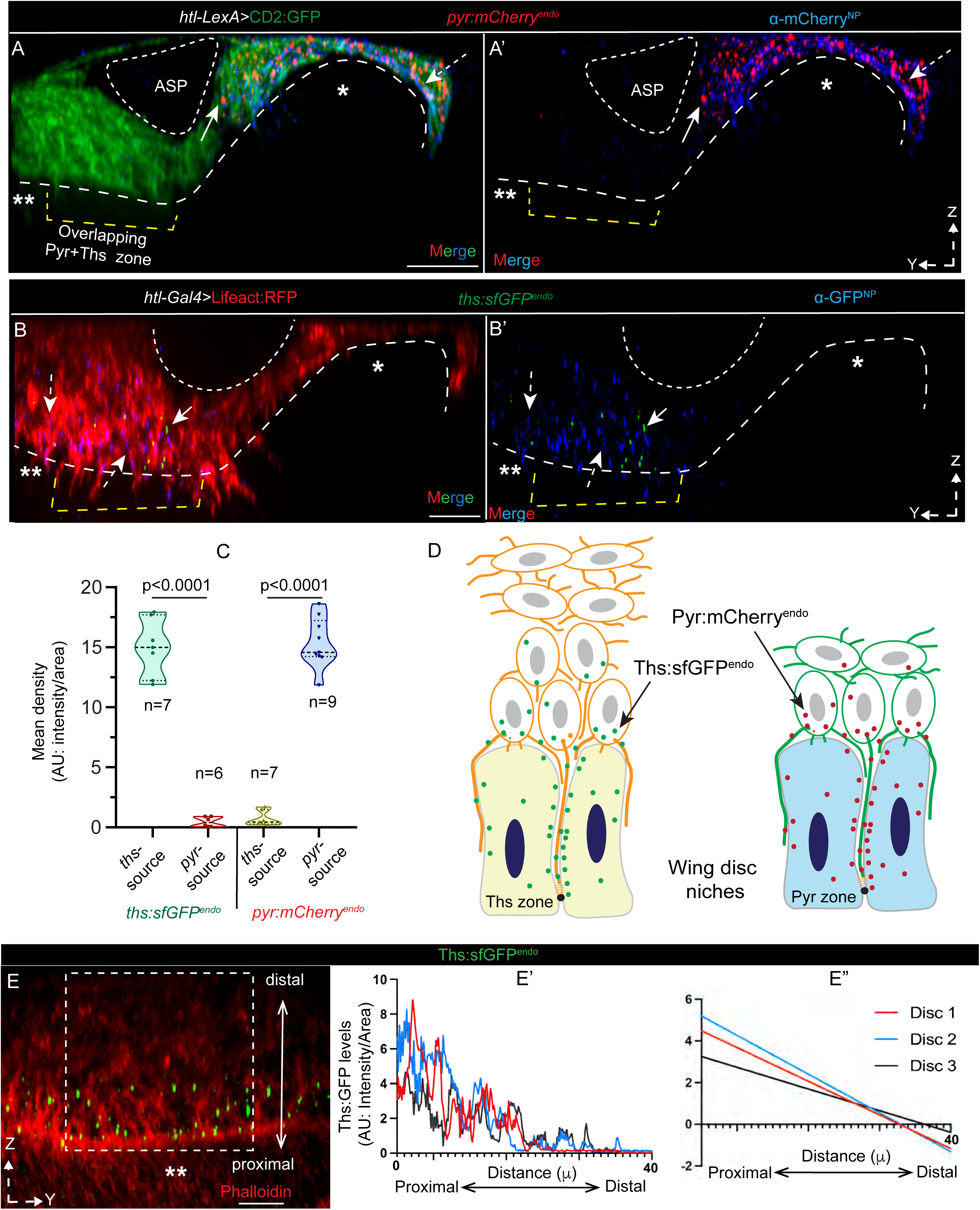
Non-overlapping niche-specific extracellular dispersion of endogenous Pyr and Ths. (A–B’) Confocal YZ views showing niche-specific, asymmetric localization of extracellular Pyr:mCherry^endo^ (α-mCherry^NP^-probed; NP-non-permeabilized condition) and Ths:sfGFP^endo^ (α-GFP^NP^-probed) on the surfaces of niche-adhering AMPs. *: *pyr*-expressing zone only; **: *ths***-**expressing zone only; overlapping zone: harbors both *pyr* and *ths* expressing cells; Arrows: intracellular ligand; Dashed arrows: extracellular ligand. (C) Quantification of Pyr:mCherry^endo^ and Ths:sfGFP^endo^ densities in AMPs (expressing CD2:GFP or Lifeact:RFP, as indicated) located in *pyr***-** or *ths***-**producing niches (n = number of wing discs; P-values from two-tailed t-tests). **(D)** Schematic summarizing results from A-C. (E-E”) Ths:sfGFP^endo^ dispersion gradient orthogonal to the *ths*-expressing zone, visualized in phalloidin-stained AMPs and wing discs. Graphs (E’,E”) show quantification of Ths:sfGFP^endo^ density along the niche-proximal to niche-distal axis (double-headed arrow) from ROIs (e.g., dashed box in E) in three independent wing discs; E”: Simple linear regression of data shown in E’ (see Materials and Methods). Genotypes and color codes are indicated in each panel. Dashed lines: AMP-wing disc boundary. Scale bar: 10 μm.

Finally, ligand dispersion patterns closely followed the niche-specific 3D morphology and AMP organization. For example, *ths*-expressing wing disc regions are populated by variable three to five orthogonal AMP layers (Figs. 2E; 3B,E). Quantification of Ths:GFP puncta density in these regions revealed that AMPs farther from the niche have progressively lower FGF reception, leading to the emergence of an asymmetric, graded signal dispersion pattern (Fig. 3E-E’’). Importantly, this high-to-low Ths gradient from niche-proximal to niche-distal layers was robustly maintained across tissues of different sizes, with the dispersion range and gradient slope scaled to AMP layer number/size (Fig. 3E-E’’). We did not detect a Pyr gradient orthogonal to the wing disc, likely because only two AMP layers overlay the *pyr*-expressing wing disc hinge. However, Pyr distribution is precisely adapted to the size and shape of these layers in different discs (Fig. 3A,A’). Together, these results reveal robust, recipient-specific, and adaptive dispersion patterns for both Pyr and Ths.

### Receptor-bound dispersion of Pyr and Ths

To determine whether the AMP-specific distribution of Ths:sfGFP^endo^ results from Htl-bound cytoneme-mediated dispersion, we first generated an *htl:mCherry^endo^* knock-in allele using CRISPR/Cas9-based genome editing (Fig.4A, Supplementary Fig. 2C, see Methods). The knock-in allele was homozygous viable and expressed a functional Htl:mCherry fusion protein at physiological levels in AMPs. As expected, Htl:mCherry localized to AMP plasma membranes. Importantly, Htl:mCherry puncta were asymmetrically enriched along the niche-adhering surfaces and cytonemes of AMPs, indicating a polarized localization of the receptor (Fig. 4B-B”).

**Figure 4.**
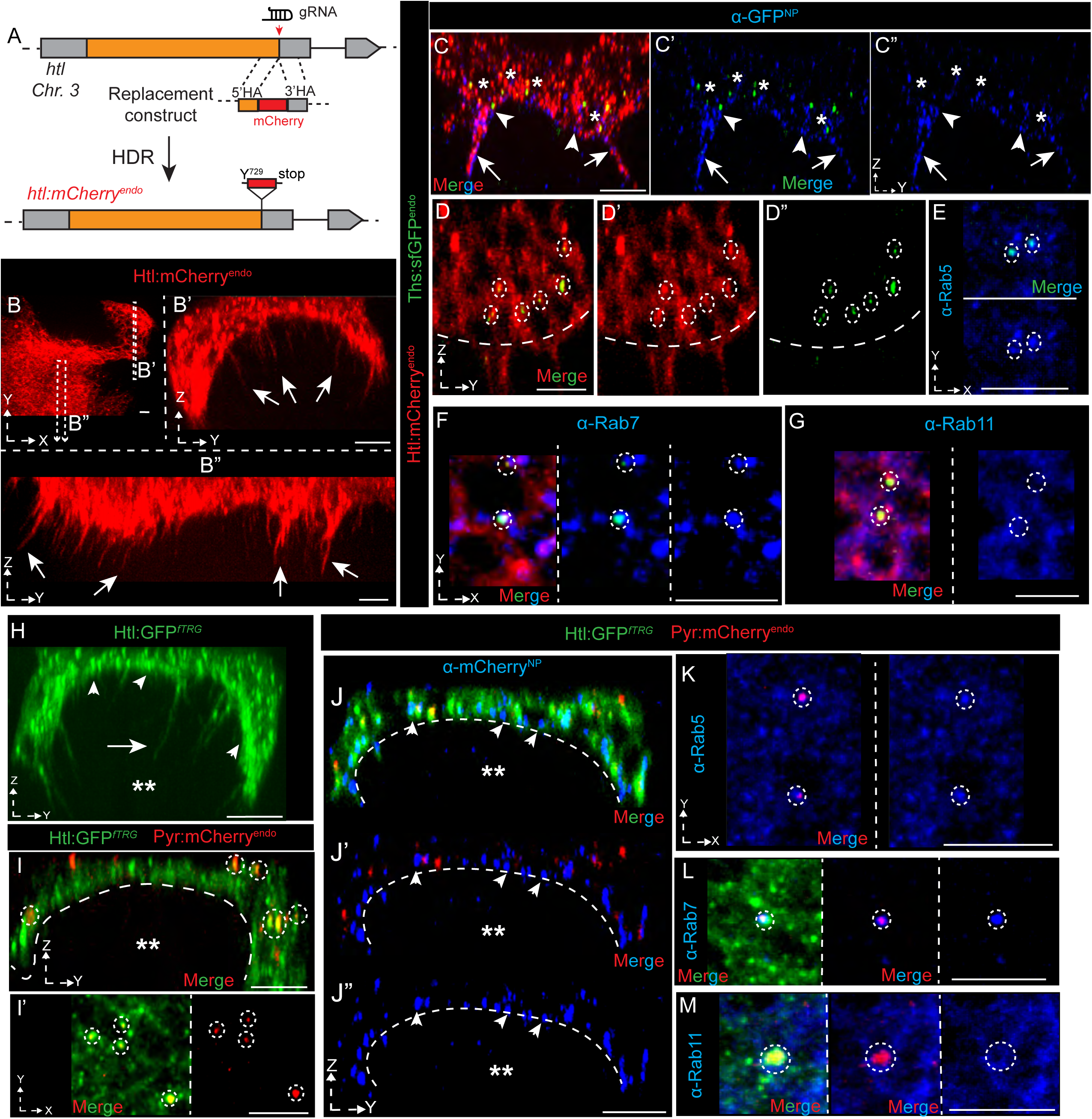
Receptor-bound extracellular segregation of Pyr and Ths. **(A)** Schematic map (not to scale) of the *htl:mCherry* genomic locus showing the genome-editing strategy and C-terminal in-frame tag insertion site (see also Supplementary Fig. 2C,D). **(B–B”)** Native Htl:mCherry expression in AMPs and localization to orthogonal cytonemes (arrows) invading the *pyr* (B,B’) and *ths* (B,B”) niches. Dashed box in B: ROI used for YZ cross-sections in B’,B”. **(C-C”)** α-GFP^NP^-stained wing discs (YZ view of the *ths* niche) from *htl:mCherry^endo^*/+, *ths:sfGFP^endo^*/+ larvae, showing enrichment of colocalized receptor-ligand puncta on the niche-proximal AMP surface, particularly on their niche-adhering sides (arrowheads) and cytonemes (arrow) of AMPs. *: nuclei of individual AMPs. **(D-G)** Intracellular co-localized ‘Htl:mCherry-Ths:sfGFP^endo^’ puncta (dashed circles) in endosomes (E-G; red channel not shown in E) only in *ths-*niche-occupied AMPs of *htl:mCherry*/+, *ths:sfGFP^endo^*/+ wing discs (single XY or YZ optical sections). Anti-Rab5, anti-Rab7, and anti-Rab11 staining marked early, late, and recycling endosomes (E-G, K-M). **(H)** Htl:GFP*^fTRG^-*expressing AMPs in the *pyr-*niche (*), showing enrichment in cytonemes (arrows) and AMP-niche junctional interface (arrowheads). **(I-M)** Co-localized ‘Htl-Pyr’ puncta from *htl:GFP^fTRG^*/+, *pyr:mCherry^endo^*/+ larvae in intracellular compartments (dashed circle; I,I’,K-M) and on the AMP surface (J-J”; probed by α-Cherry^NP^), particularly enriched at the niche-proximal side (arrowhead; J-J”) of the AMP surface; **: *pyr-*niche. Anti-Rab5, anti-Rab7, and anti-Rab11 staining (K-M). Genotypes, merged/split channels, and color codes are indicated in each panel. Also see Supplementary Figure 3 for quantitative data, and Supplementary Figure 4 for comparison of extracellular Ths:GFP^endo^ in niche-proximal and niche-distal AMPs. Dashed lines: AMP-wing disc boundary. Scale bars: 10 µm.

Non-permeabilized α-GFP^NP^ staining of wing discs from *ths:sfGFP^endo^/+; htl:mCherry^endo^/+* wing discs revealed that externalized Ths:sfGFP^endo^ precisely colocalized with Htl:mCherry puncta on AMP and cytonemes surfaces (Fig.4C-C”). Notably, α-GFP^NP^-probed Ths:sfGFP^endo^ was concentrated on AMP surfaces facing the wing disc *ths* source, paralleling Htl:mCherry distribution, consistent with receptor-dependent, recipient-specific ligand transfer (Fig. 4C-C’’). In addition, bright GFP-positive Ths:sfGFP^endo^ puncta not probed by aGFP^NP^ represented internalized ligand pools, which were also colocalized with Htl:mCherry.

To characterize the localization of this internalized pool of receptor-bound Ths:sfGFP^endo^ puncta in the early, late, and recycling endosomal compartments, we immunostained *ths:sfGFP^endo^/+; htl:mCherry^endo^/+* wing discs with α-Rab5, α-Rab7, and α-Rab11 antibodies. These experiments showed that the majority of internalized Ths:sfGFP^endo^::Htl:mCherry puncta were in early (Rab5-positive; 85.3 ± 2.2%) or late (Rab7-positive; 80.4 ± 4.9%) endosomes, but absent from recycling (Rab11-positive) endosomes (Figs. 4D-G; S3A).

We next examined Htl-bound Pyr:mCherry^endo^ distribution using the *htl:GFP* ^fTRG^ fly line (Sarov et al., 2016), which expresses functional, cytoneme-localized Htl:GFP at physiological levels (Fig 4H and (Patel et al., 2022)). In *pyr:mCherry^endo^*/+; *htl:GFP* ^fTRG^ /+ wing discs, 100% of Pyr:mCherry^endo^ puncta (n = 119 from three discs) outside the source colocalized with Htl:GFP (Fig.4I,I’). Non-permeabilized α-mCherry^NP^ staining confirmed precise colocalization of Pyr:mCherry^endo^ with Htl:GFP on AMP surfaces, with asymmetric enrichment at the niche::AMP interface (Fig. 4J-J’’’). These results suggest Htl-dependent non-cell-autonomous Pyr transfer.

Similar to Ths:sfGFP^endo^, intracellular Pyr:mCherry^endo^ puncta colocalized with Htl:GFP in early (Rab5-positive; 68.9 ± 7.7%; n = 77) and late (Rab7-positive; 86.2 ± 2.4%; n = 106) endosomes, but not in recycling (Rab11-positive) endosomes (Figs.4K-M; S3A). Together, these results show receptor-bound distribution of both Pyr and Ths ligands, explaining their precisely recipient-specific dispersion patterns. These results further suggest that the externalized ligand pool remains bound to Htl on AMP surfaces prior to receptor-mediated endocytosis. The absence of Htl-bound ligands from recycling endosomes likely reflects endosomal signaling and/or trafficking toward termination of FGF signaling.

### AMP cytonemes are required for segregated Pyr and Ths dispersion

Building on our previous demonstration of cytoneme-mediated exchange of overexpressed Pyr:GFP and Ths:GFP (Patel et al., 2022), and the generation of *pyr:mCherry^endo^* and *ths:sfGFP^endo^* knock-in construct, we sought to genetically dissect the cytoneme-driven mechanisms underlying the asymmetric distribution patterns of endogenous FGFs. However, tracking FGFs in live tissue proved challenging due to the low fluorescence intensity of the endogenous FGF chimeras and the sub-150 nm diameter of orthogonal AMP cytonemes, which dynamically invade through the ∼80 μm-thick wing disc epithelium. To overcome these limitations, we employed super-resolution Airyscan imaging under fixed-tissue conditions (suitable for low-light imaging). In addition, we used non-permeabilized immunostaining to detect externalized Ths:sfGFP^endo^ (with α-GFP^NP^) and Pyr:mCherry^endo^ (with α-mCherry^NP^). As shown in Figures 5A-E’, cytonemes were visualized by expressing CD2:GFP, Lifeact:RFP (F-actin probe), or by phalloidin staining (F-actin probe), following established methods (Patel et al., 2022). These combined approaches substantially improved detection of Pyr:mCherry^endo^ (Fig. 5A) and Ths:sfGFP^endo^ along cytonemes (Fig. 5B-E’).

**Figure 5.**
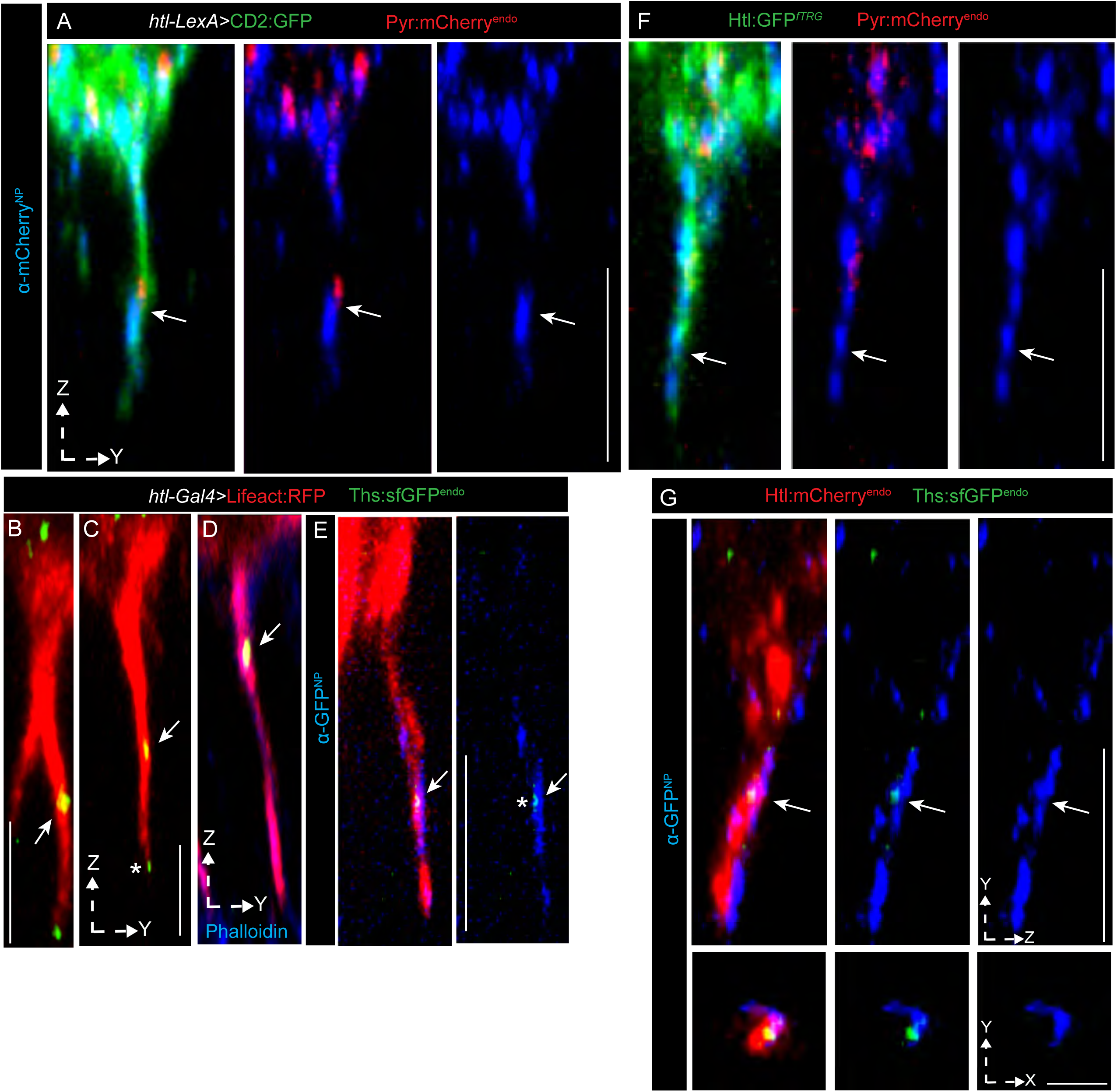
Endogenous Pyr and Ths localize to Htl-containing AMP cytonemes. **(A-E)** High-resolution images of orthogonal AMP cytonemes marked by CD2:GFP (A) or Lifeact:RFP (B-E), showing Pyr:mCherry^endo^ (A) and Ths:sfGFP^endo^ (B-E) localization (arrows) along cytonemes (extended YZ projection from 1-5 optical sections shown; B-D: Airyscan super-resolution). α-mCherry^NP^ (A) and α-GFP^NP^ (E) detects Pyr:mCherry^endo^ and Ths:sfGFP^endo^ on cytoneme surfaces, respectively. **(F-G)** Extended YZ projection from 1-5 optical sections of orthogonal AMP cytonemes with colocalized Htl:GFP*^ftRG^*-Pyr:mCherry^endo^ (F) or Htl:Cherry-Ths:GFP^endo^ (G) puncta along cytonemes. α-mCherry^NP^ (F) and α-GFP^NP^ (G) detects Pyr:mCherry^endo^ and Ths:sfGFP^endo^ along cytoneme surfaces, respectively. Lower panels in G: XY cross sections at the arrow-indicated position from corresponding upper panels, showing co-localization of Ths with Htl precisely on cytonemes. Genotypes: (A) *pyr:mCherry^endo^; htlLexA, LexO-CD2:GFP*, (F) *pyr:mCherry^endo^/+; htl:GFP^fTRG^/+,* (B-E) *ths:sfGFP^endo^; htlGal4, UAS-Lifeact:RFP*, (G) *ths:sfGFP^endo^ /+; htl:mCherry^endo^/+*. Merged/split channels and color codes are indicated in each panel. Also see Supplementary Figure 4B,C for quantitative analyses of cytoneme localization and controls for antibody-probed FGF localization on AMP/cytoneme surfaces. Scale bars: 10µm; 1µm for G lower panels.

Almost all cytonemes from niche-proximal AMPs invaded through the intercellular junctions of the wing disc epithelium and received FGF ligands on their surfaces (Figs. 5B-E’;S4A-A”’). In contrast, AMPs located in the distal-most layers away from the wing discs lacked detectable cell-surface FGFs (Fig.S4A). Examination of wing discs co-expressing either Htl:GFP*^fTRG^* with Pyr:mCherry^endo^ or Htl:Cherry with Ths:GFP^endo^, revealed colocalization of native receptors and ligands on AMP cytoneme surface (Fig.5F,G). Nearly all analyzed Htl-containing cytonemes invading the wing disc epithelium displayed co-localized ligands (Figs. 5F,G; S4B). To confirm that receptor- and cytoneme-dependent exchange is required for the non-cell-autonomous dispersion of both FGFs, we knocked down *htl* in AMPs by expressing *htl* RNAi under *htl*-*Gal4* in the *ths:sfGFP^endo^* and *pyr:mCherry^endo^* flies (Fig.S4C-E). Consistent with previous findings (Patel et al., 2022), loss of Htl suppressed orthogonal cytoneme formation, abolishing niche occupancy, FGF signaling, and AMP proliferation. Under these conditions, both extracellular (probed by α-GFP^NP^ or α-Cherry^NP^) and intracellular ligand pools (probed by GFP or mCherry fluorescence) were completely absent (Fig.S4C-E), confirming that AMP-specific asymmetric ligand dispersion patterns depend on both Htl and cytonemes from AMPs.

### Spatial regulation of cytoneme polarity and specificity

Because Ths:sfGFP^endo^ disperses over a more extended range than Pyr, and cytoneme-deficient clones can be more frequently detectable within the multi-stratified AMP layers over the *ths* niche than the *pyr* niche (Patel et al., 2022), we used mosaic analyses in the *ths* niche to directly examine the polarity of cytonemes across AMP layers. We generated sparsely located CD8:RFP-marked AMP clones (*htl>FRT>Gal4*) expressing *diaphanous* (*dia*; formin) *RNAi* in *ths:sfGFP^endo^* larvae. Expression of *dia* RNAi has been validated across multiple cell types, including AMPs, as a means to disrupt cytoneme formation without impairing intrinsic ligand-binding capacity or downstream signal transduction (Patel et al., 2022; Roy et al., 2014).

In wild-type controls, CD8:RFP-marked clones were distributed across all niche-proximal and niche-distal layers. Moreover, in this condition, both unmarked and marked niche-proximal AMP clones readily received Ths:sfGFP^endo^, indicating that the expression of CD8:RFP did not impair Ths uptake in AMPs (Fig. 6A-C). In contrast, cytoneme-deficient *dia* RNAi clones exited the niche, localized distally, and failed to take up Ths:sfGFP^endo^ (Fig. 6C-E’). This absence of ligand uptake in *dia-*deficient clones parallels the loss of pMAPK signaling previously reported in *dia*-mutant AMPs (Patel et al., 2022). These results indicate that AMP cytonemes are required for establishing graded Ths:sfGFP^endo^ dispersion patterns.

**Figure 6.**
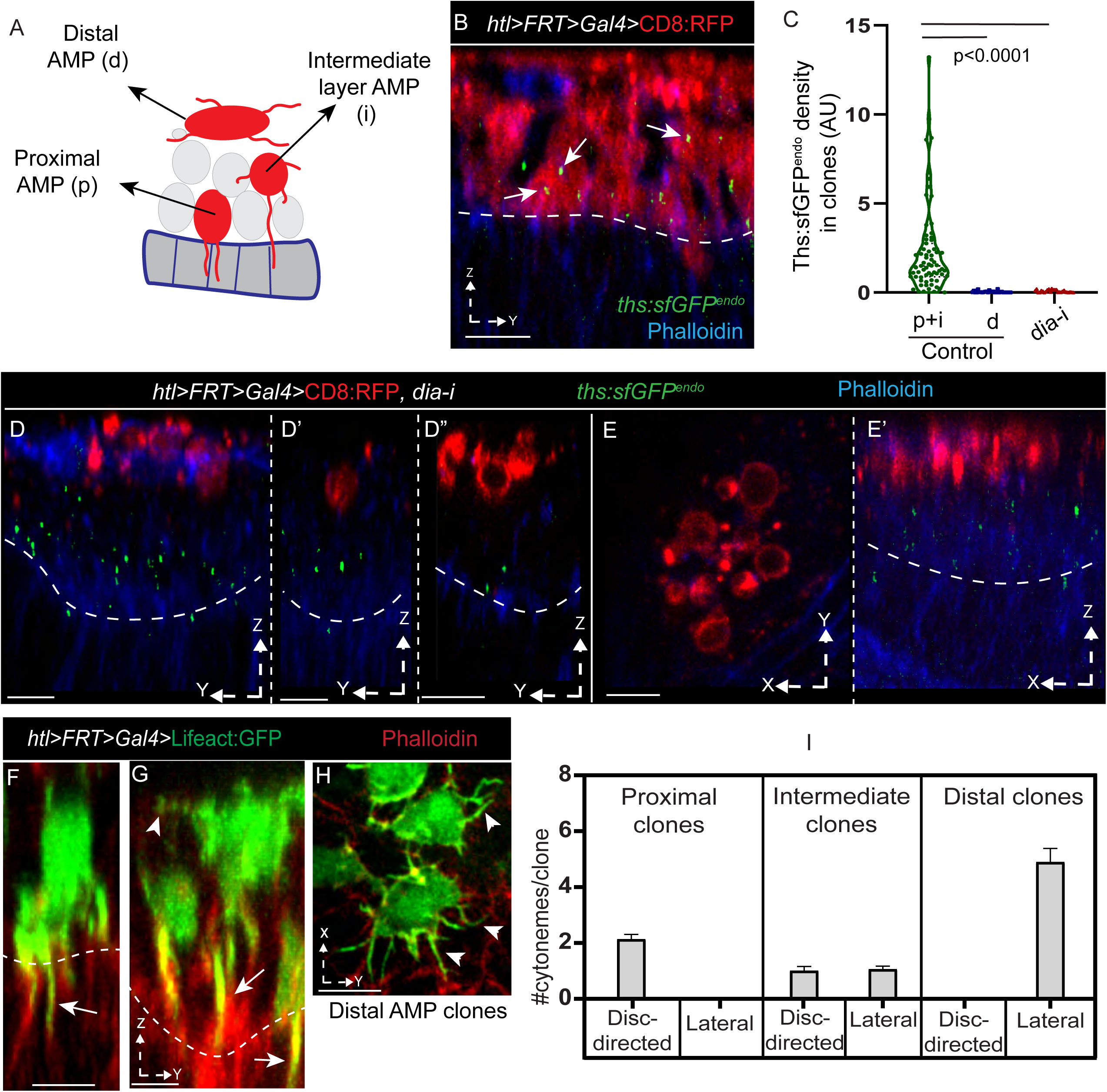
Position-dependent abundance of orthogonal AMP cytonemes underlies Ths dispersion patterns. **(A-E’)** Sparsely localized CD8:RFP-marked AMP clones expressing control (B; *hsflp/+; UAS-CD8RFP/+*) or *dia*-*RNAi* (D-E’; *hsflp/+; UAS-CD8RFP/+; htl>FRT>Gal4/UAS-dia-RNAi*,) in *ths*:*sfGFP^endo^* wing discs. A: Schematic, showing AMP clones in different cell layers scored for analyses. Control WT AMP clones in niche-proximal layers take up Ths:GFP^endo^ (arrow), whereas *dia*-RNAi clones localize distally and lack Ths:GFP^endo^ uptake (D-E’). C: Graphs comparing Ths:sfGFP^endo^ densities in AMPs in WT (in both proximal+intermediate and distal AMPs) and *dia* deficient AMPs. P value-ANOVA with Tukey HSD (HSD[0.05]=14.91). N = WT: 67 proximal +intermediate clones and 28 distal clones (6 wing discs); *dia-i* mutant: 5 wing discs, 25 clones. **(F-I)** High-resolution images of sparse CD8:GFP-marked AMP clones in *ths* niche (*hsflp/+; UAS-Lifeact:GFP/+; htl>FRT>Gal4/+*), showing orthogonal (arrow) and lateral (arrowhead) cytonemes from niche-proximal and niche-distal AMPs, respectively. I: Graphs comparing the numbers of cytonemes and their niche-specific polarity from different orthogonal positions relative to the wing disc niche (also see Supplementary Figure 5A for numbers). Disc orientations, merged/split channels, color codes, and genotypes are indicated in each panel. Dashed lines: AMP-niche interface. Scale bars: 10µm.

Our previous work (Patel et al., 2022) showed that the niche-specific orthogonal polarity of cytonemes is confined to niche-adhered AMPs, whereas niche-distal AMPs extend only lateral cytonemes. This led to a hypothesis that the AMPs at different orthogonal distances from the niche produce varying numbers of niche-directed cytonemes, and that Ths/FGF reception via these cytonemes leads to the emergence of precise, recipient-specific shapes of ligand dispersion gradient. To examine this possibility, we generated and examined small-sized (1-2 cell) AMP clones marked by Lifeact:GFP and quantified cytoneme number and polarity from AMP clones located at different distances away from the wing disc niche.

As expected, single-cell clones immediately adjacent to the niche extended only orthogonal cytonemes (Fig. 6F; Fig. S5A). However, clones in the second, third, and fourth orthogonal layers projected progressively fewer orthogonal cytonemes, accompanied by a corresponding increase in short, randomly oriented lateral cytonemes (Figs. 6G-I; Fig. S5A). These findings reveal a gradual decline in FGF-receiving orthogonal cytoneme number with increasing niche distance, supporting the model in which signal-specific cytoneme polarity and cytoneme-mediated signal reception/niche adhesion are spatially regulated in a location-dependent manner.

### Feedback from the Htl-Pnt signaling axis sustains ligand-specific polarity of AMP cytonemes

The cell-autonomous requirement for Htl in AMPs to generate orthogonal, niche-directed cytonemes suggested that FGF signaling may provide feedback to regulate niche (or signal)-specific cytoneme polarity, and, thereby, self-reinforce the niche-specific positional information. To test whether the intracellular Htl pathway contributes to this feedback, we examined the ETS transcription factor Pointed (Pnt), a downstream effector of Htl signaling shown in *Drosophila* blood progenitors (Dragojlovic-Munther and Martinez-Agosto, 2013). To visualize native Pnt expression pattern, we used a recombineered Pnt:GFP BAC transgene harboring fly line (Venken et al., 2009), which expresses the functional protein under its native genomic regulatory elements, and compared its expression to dpERK staining, a pMAPK readout of FGF signaling (Fig.7A-D).

**Figure 7.**
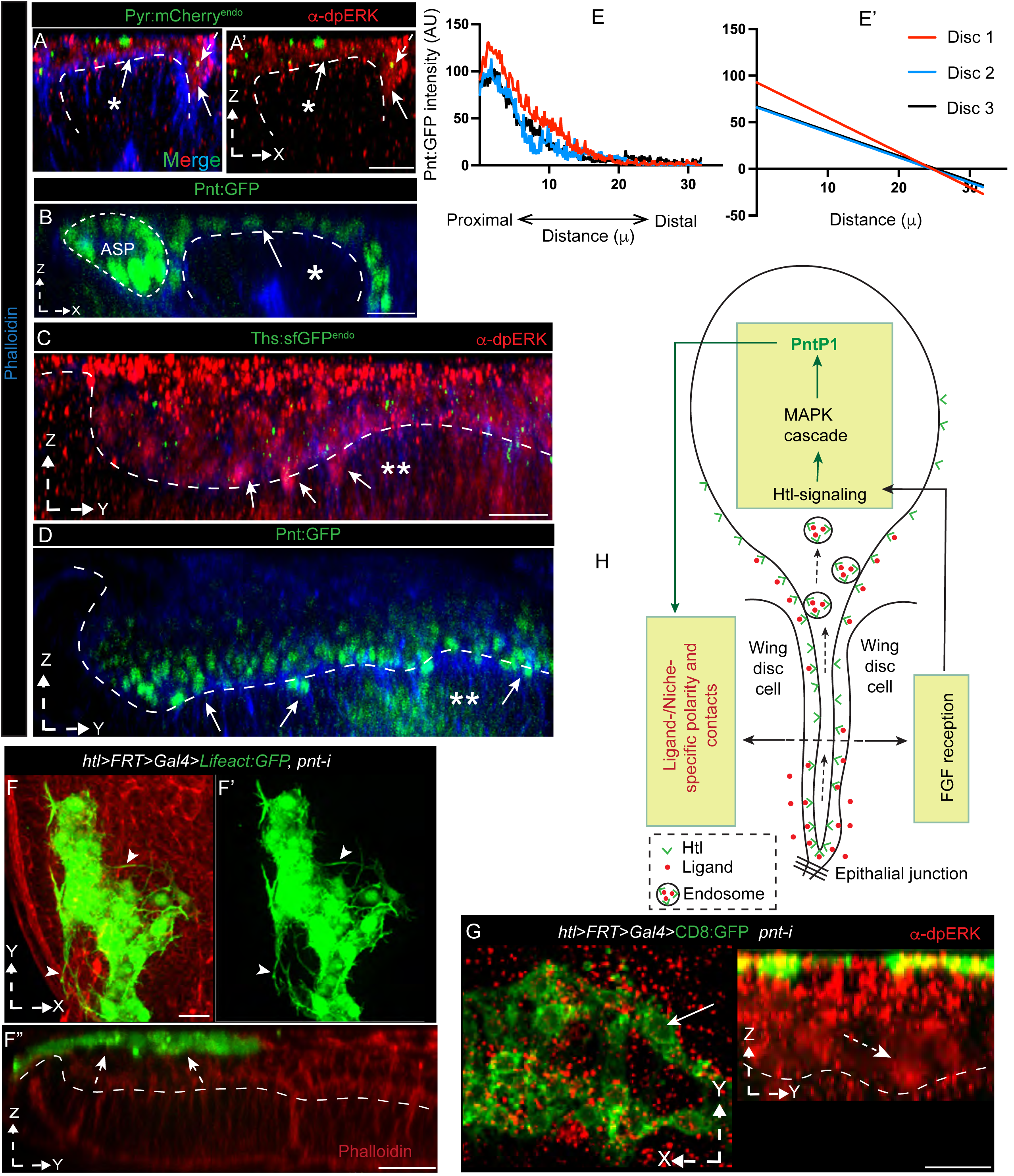
Htl-Pointed signaling feedback sustains orthogonal FGF-receiving AMP cytonemes. **(A–D)** Endogenous Pnt:GFP expression (B,B’,D; arrow) parallels Pyr:mCherry^endo^ and Ths:sfGFP^endo^ reception and dpERK activation (pMAPK; A,A’,C; arrow) in niche-proximal AMPs, with graded (high-to-low) nuclear intensity from proximal to distal layers in both *pyr* (*) and *ths* (**) niches. **(E,E’)** Quantification of Pnt:GFP gradient along orthogonal AMP axis in three independent *ths*:sfGFP^endo^ wing discs (E’-simple linear regression of E). **(F-F”)** *pnt*-RNAi-expressing AMP clones (CD8:GFP-marked; *hsflp/+; UAS-Lifeact:GFP/+; htl>FRT>Gal4/UAS-pnt-RNAi*, (H,H’) *hsflp/+*) exit the niche, localize distally, lose orthogonal cytonemes (dashed arrow), but retain lateral cytonemes (arrowhead) and, apparently, gain fusogenic responses. Also see Supplementary Figure 5A,B for quantitative data. **(G)** dpERK staining (dashed arrow) in wing discs harboring *pnt*-RNAi-expressing AMP clones shows the absence (arrow) of pMAPK activation in *pnt*-RNAi-expressing clones (*hsflp/+; UAS-CD8:GFP/+; htl>FRT>Gal4/UAS-pnt-RNAi*). Also see Supplementary Figure 5C for quantitative data. **(H)** Model: AMP-intrinsic Htl-Pnt feedback sustains ligand- and niche-specific orthogonal cytonemes and thereby self-sustaining specific FGF reception via cytonemes. Disc orientations, merged/split channels, color codes, and genotypes are indicated in each panel. Scale bars: 10µm.

In *ths:sfGFP^endo^* and *pyr:mCherry^endo^*wing discs, all ligand-receiving niche-proximal AMPs showed nuclear dpERK, indicating activation of FGF signaling. Similarly, Pnt:GFP was localized to the nuclei of niche-adhering AMPs, in both *ths*- and *pyr*-producing niches (Fig. 7B,B’,D), consistent with its role as a transcription effector, downstream of pMAPK signaling. Notably, nuclear Pnt:GFP displayed a high-to-low gradient from niche-proximal to niche-distal AMPs, paralleling dpERK intensity (Fig. 7A-D). Quantitative analysis across multiple independent wing discs confirmed that this Pnt:GFP gradient was robust and scaled with AMP layer size and organization (Fig. 7E,E’). The graded patterns of nuclear-localized Pnt, in parallel with the graded Ths:sfGFP^endo^ reception and pMAPK signaling, indicate a spatially asymmetric activation of Htl-Pnt signaling axis (Figs. 3E,E’;7A-E’).

To determine whether Pnt can feedback regulate niche-specific affinity, orientation, and activity of AMP cytonemes, we generated *pnt* gain-of-function (GOF: *Hs-Flp*; *htl>FRT>Gal4* driving *UAS-PntP1*) and loss-of-function (LOF: *Hs-Flp; htl>FRT>Gal4* driving *UAS-pnt-RNAi*) AMP clones (see Materials and Methods). Despite multiple attempts, CD8:GFP-marked *Pnt*P1 GOF clones (similar to *htl* GOF clones) were not recovered from >120 wing discs, likely due to lethality caused by overexpression of the *pnt*, indicating critical functions of the gene in AMPs. In contrast, we successfully obtained wing discs with CD8:GFP-marked AMP clones expressing *pnt*-*RNAi* (Fig. 7F-F”).

Unlike control clones, which populated all orthogonal AMP layers (Figs. 6A-C), *pnt*-RNAi-expressing clones (*n* = 52) exited the niche and localized exclusively to niche-distal layers, losing niche-specific orthogonal polarity and niche-adhering cytonemes, but adopting the morphological features of distal *WT* AMP clones, aligning parallel to the wing disc plane and retaining lateral cytonemes (Figs. 6I; 7F-F””; S5A,B). These clones also adhered to each other, mimicking a fusogenic response previously observed in *htl-*deficient AMP clones (Patel et al., 2022). Furthermore, dpERK staining revealed that *pnt*-deficient AMP clones lacked detectable pMAPK signaling (Figs. 7G; S5C). These *pnt* LOF phenotypes closely resembled those of *htl*-RNAi-expressing AMP clones, supporting a shared cell-autonomous requirement of both Htl and Pnt in maintaining orthogonal cytoneme polarity and niche occupancy. These results establish the Htl-Pointed feedback loop as a key intrinsic auto-regulator of AMP cytoneme polarity, ensuring recipient-specific Pyr and Ths dispersion and niche-directed patterning.

## DISCUSSION

### Non-overlapping dispersion of multiple ligands within a shared receptor microenvironment

How a small set of evolutionarily conserved signaling pathways generates information for diverse, context-specific morphogenetic responses remains a long-standing question. This study provides insights into how the spatial segregation of multiple ligand dispersion, maintained through cytoneme-mediated transport and signaling feedback, enables conserved pathways to generate and amplify differential information. We directly visualize the native, non-overlapping dispersion of Pyr and Ths, demonstrating that their segregation is maintained and scaled by polarized, cytoneme-mediated, Htl-bound transport, coupled with signaling feedback regulation of cytoneme polarity and specificity. Using high-resolution imaging of fully functional, endogenously tagged knock-in Pyr, Ths, and Htl constructs, together with immunoprobing of secreted ligands and labeling of AMP and wing disc membranes, we could precisely map the extracellular and intracellular distribution of Pyr and Ths.

Despite being secreted FGF8 family proteins, capable of long-range diffusion (Harish et al., 2023; Toyoda et al., 2010), and despite their juxtaposed sources and shared use of the Htl receptor in isogenic AMPs, Pyr and Ths exhibited extracellular distributions that were strikingly local, restricted only to niche-adhering AMPs, maintaining robustly non-overlapping and AMP-specific distribution. Quantitative analyses further suggested position-dependent graded reception of FGFs among orthogonally organized AMPs, and scaling of their dispersion range to tissue size and number of orthogonal AMP layers. Overlapping patterns of ligand, receptor, and downstream signaling activation indicated tight spatial coordination between signal uptake and signaling response. These results collectively establish an adaptive, precise, and robustly recipient- and niche-specific non-overlapping dispersion of Pyr and Ths within the niche-occupied AMPs.

### Cytoneme polarity and specificity are tuned in a position-dependent manner within the AMP niche

Our results show that the segregated AMP-specific patterns of Pyr and Ths dispersion are a consequence of cytoneme-mediated Htl-bound dispersion. Loss of either cytonemes (via *dia* knockdown) or Htl (via *htl* knockdown) abolished ligand reception and eliminated extracellular ligand pools, indicating that Pyr and Ths are not freely released, and/or pre-existing in the extracellular space. Instead, they are exchanged directly at cytoneme-source contact points only when cytonemes occupy the intercellular space of the FGF-producing disc epithelium. Analyses of extracellular and endosomal localization suggest that Htl on the niche-occupying cytonemes surface first receives Pyr and Ths directly by adhering to the intercellular junction of the FGF-source epithelium. Subsequently, receptor-mediated endocytosis internalizes ligands within the AMP cell body, preserving precise, AMP-specific niche-restricted distribution.

Significantly, the number and polarity of these FGF-receiving cytonemes vary with AMP position. Niche-adhered AMPs project numerous orthogonal cytonemes traversing the niche epithelium, while niche-distal AMPs lack them and instead project shorter, randomly oriented lateral cytonemes. This graded polarity of FGF-receiving cytonemes correlates with the levels of ligand reception, providing a structural explanation for the emergence of AMP-specific signal and signaling gradients. Together, these results suggest that position-dependent tuning of cytoneme polarity and specificity provides a scalable mechanism to generate ligand dispersion matching the tissue size and AMP organization.

### Autoregulatory signaling feedback reinforces ligand-specific cytoneme polarity

A central finding of this study is the discovery of a cell-intrinsic positive feedback loop downstream of Htl signaling that reinforces ligand-specific cytoneme polarity. We show that the ETS transcription factor Pnt is asymmetrically activated in a high-to-low gradient from niche-proximal to distal AMPs, mirroring dpERK activity and ligand uptake pattern. Secondly, loss of Pnt in AMP clones abolishes orthogonal cytoneme formation and pMAPK signaling, causing these clones to exit the niche and acquire fusogenic behaviors, all of which phenocopy *htl* loss-of-function AMP clones(Patel et al., 2022). Notably, similar to *htl-*deficient AMP clones, *pnt-*deficient AMP clones retain lateral cytonemes, while exclusively lacking orthogonal FGF-receiving cytonemes. These results suggest that a shared Htl-Pnt pathway constitutes a dedicated feedback module that can specifically maintain niche-directed, ligand-specific orthogonal polarity of cytonemes. This finding establishes cytoneme-mediated signaling as an autoregulated, self-organizing process and provides insights into the molecular components involved in eliciting the feedback.

A key consequence of this positive feedback loop is that it places the polarity of FGF-receiving cytonemes under the direct control of Htl signaling pathway, driven by the very ligand transported by Htl-containing cytonemes within each AMP. By coupling contact-mediated ligand reception to feedback-driven cytoneme polarity/affinity, the same receptor-pathway module can extract distinct positional and fate information depending on whether AMPs adhere to Pyr- or Ths-expressing cells. This mechanism explains how Pyr and Ths, acting through Htl, segregate in space during their dispersion and promote divergent AMP lineages and organizations. These results also demonstrate that cytoneme polarity and specificity are actively regulated by a cell-intrinsic feedback mechanism, enabling cytonemes to concurrently sense, respond to, and self-adjust their specificity for each ligand and ligand source, even when using the same receptor.

Previously, we showed that Pnt is also a downstream effector of the Drosophila FGFR Breathless (Btl) in the ASP, where the Btl-Pnt axis feedback regulates the polarity and number of Bnl-receiving cytonemes to scale recipient-specific Bnl gradients and signaling patterns(Du et al., 2018a). The discovery of similar Pnt-dependent feedback in different cell types, and also for multiple ligands converging on the same receptor, suggests that signaling feedback is a generalizable principle for sustaining cytoneme polarity and specificity. Intriguingly, this simple autoregulatory principle governing cytoneme polarity and signaling specificity could also operate when multiple receptors converge on the same downstream pathway. Thus, such simple signaling feedback-driven autoregulation can make cytonemes versatile organizers of complex signaling microenvironments.

Several open questions remain. We do not know the molecular effectors linking Pnt activity to cytoskeletal polarization of cytonemes or whether additional feedback architectures operate in the AMP niche to refine ligand territories. Although technically challenging to track the dynamics of endogenous ligands in the deep tissue AMP cytonemes, future improved tools for the live imaging of endogenous ligand-receptor complexes will help uncover the ligand-specific temporal dynamics of cytoneme turnover and feedback regulation. Htl signaling plays essential roles in embryonic myogenesis, cardioblast fusion, visceral muscle development, hematopoiesis, and neuro-glia interactions (Dragojlovic-Munther and Martinez-Agosto, 2013; Franzdóttir et al., 2009; Irizarry and Stathopoulos, 2015; Macabenta and Stathopoulos, 2019; Kadam et al., 2009a; McMahon et al., 2010; Reim et al., 2012; Rothenbusch-Fender et al., 2017; Stork et al., 2014; Wu et al., 2017; Patel et al., 2022; Dutta et al., 2005; Vishal et al., 2020). It would be interesting to explore whether autoregulated cytoneme-mediated Pyr and Ths segregation underlies these morphogenetic events. Finally, testing how the same principle for cytonemes is programmed and used by other signaling pathways, stem cell niches, and developmental fields to organize multiple signaling cues would reconcile how distinct patterning outcomes arise using cytoneme-mediated signaling.

## MATERIALS AND METHODS

### Drosophila melanogaster strains

All flies were raised at standard conditions (Patel et al., 2022). The list of flies used and generated during this study is provided in Supplementary Table 1.

### Molecular cloning and generation of transgenic *Drosophila*

DNA cloning experiments followed standard molecular biology protocols (Patel et al., 2022). Molecular biology reagents, including PCR primers, kits, and DNA cloning methods are listed in Supplementary Table 1. DNA constructs were sequence-verified prior to use in the generation of transgenic flies via P-element-mediated germline transformation (performed by Rainbow Transgenic Flies, Inc.).

### CRISPR/Cas9-based genome editing for generating *pyr:mCherry^endo^, ths:sfGFP^endo^ and htl:mCherry^endo^* knock-in alleles

We generated *pyr:mCherry^endo^, ths:sfGFP^endo^*and *htl:mCherry^endo^* knock-in constructs following a standard method described earlier (Du et al., 2018b; a, 2017; Gratz et al., 2015). Tag insertion sites were chosen to be identical to the previously reported for *UAS-Pyr:GFP2, UAS-Ths:GFP2* and *LexO-Htl:mCherry* constructs (Patel et al., 2022). Single gRNA target site was selected within the fifth coding exon of *pyr* and *ths* and first coding exon for *htl* using the tools described in (Du et al., 2018a). Supplementary Table 1 indicates the gRNAs (cloned in pCFD3) with zero off-targets (PAM is underlined) used for targeting *pyr*, *ths*, and *htl*, and methods for generating replacement cassettes: *pJet-pyr:mCherry^endo^*, *pJet-ths:sfGFP^endo^*, and *pUC19-htl:mCherry^endo^*. The set of specific gRNA and replacement cassette per gene was injected into the *{nos-Cas9}ZH-2A* (BL 54591) *e*mbryos, and subsequent progenies were screened and sequence-verified for ends-out HDR following (Du et al., 2018a). In Pyr:mCherry^endo^, mCherry was inserted between “…TTTT_208_” and “T_209_PAS…”. In Ths:sfGFP^endo^, sfGFP was inserted between “…PKKS_137_” and “V_138_NDA…”. In Htl:mCherry, mCherry was inserted at the C-terminus, between the last amino acid and stop codon of the *htl* gene (see Figure 4A). The strategy and results of the HDR screen are summarized in Supplementary Figure 2A-D).

### Immunohistochemistry

All immunostainings, including detergent-free EIF-staining against GFP and mCherry, were carried out following protocols described in (Du et al., 2018a). Primary and secondary antibodies and their dilutions are in Supplementary Table 1.

### Mosaic analyses

To generate FLP-out clones of AMPs of various genotypes, females of *hsflp; htl>FRT>stop>FRT>Gal4; UAS-X* flies (*X=UAS-CD8:GFP, UAS-CD8:RFP* or *UAS-Lifeact:GFP)* were crossed to *w^1118^* (control), *UAS*-*diaRNAi* (for *dia* knockdown), or *UAS*-*htlRNAi* (for *htl* knockdown) or *UAS*-*pntRNAi* (for *pnt* knockdown) male flies. To generate FLP-out clones of AMPs for analyzing uptake of Ths:sfGFP^endo^, females of *hsflp; ths:sfGFP^endo^; htl>FRT>stop>FRT>Gal4* flies were crossed to *UAS-mCherryCAAX* (control) or *UAS-mCherryCAAX; UAS-diaRNAi* (test). Crosses were reared at room temperature and clones were induced by following (Patel et al., 2022).

### Lethality rescue by transgenic and genome-edited constructs

*DfBSC25/CyO* (*pyr* and *ths* null), *Ths^759^/CyO* (*ths* mutant), *ths-Gal4/CyO* (*ths* null), and *pyr-Gal4/CyO* (*pyr* null) flies were used as null alleles. 5 males from each of these mutant lines were crossed with 5 females of the test fly line. 3 such crosses were set up for each mutant-test line pair. Fly lines tested for their ability to rescue mutants were: *UAS-Pyr:GFP1, UAS-Pyr:GFP2, UAS-Ths:GFP1, UAS-Ths:GFP2, pyr:mCherry^endo^*, and *ths:sfGFP^endo^*. For *UAS-X* constructs, “*DfBSC25 (or Ths^759^*) */CyO; UAS-X”* was crossed to specific *fgf*-*Gal4* (null alleles). Surviving Non-*CyO* F1 flies were counted and their emergence was considered as complete rescue; data provided in “Source data Fig.S2B”. *pyr:mCherry^endo^* (M1-10, M1-14) and *ths:sfGFP^endo^* (M9-2, M18-6, F9-2) were used for this and all other experiments.

### Egg-deposition assay

To determine the number of legs laid by a female in 24 hours, one 7-day old female fly of each genotype (1: *ths-Gal4/+*; 2: *ths-Gal4/UAS-Ths:GFP1*; 3: *ths-Gal4/UAS-Ths:GFP2*) was crossed with two 7-day old *w1118* male flies. 10 such crosses for each genotype were set up. The eggs were collected on apple juice plates, and number of eggs deposited on each plate was counted after 24 hours of setting up the cross.

### Protein structure analysis

PDB files containing the predicted structures for Pyramus (AF-B9ZW35-F1-v4) and Thisbe (AF-Q6Q7I9-F1-v4) proteins were obtained from AlphaFold Protein Structure Database (https://alphafold.ebi.ac.uk/) and analyzed using ChimeraX 1.9 software (see Supplementary Table 1).

### Microscopy and image processing

We used a Yokogawa CSUX1 spinning disc confocal system equipped with an iXon 897 EMCCD camera, Zeiss LSM 900 with Airyscan 2 and GaAsP detectors, and Leica Stellaris 8 microscope with HydS and HydX detectors. Images were resolved using either 20X (0.5NA), 40X (1.3NA oil), 60X (1.4NA oil, Spinning disc), or 63X (1.4NA oil for LSM Airyscan) objectives. Bright filed images of adult ovaries were taken using Zeiss Discovery V8 SteREO microscope with AxioCam HRc camera. All images were processed using Fiji/ImageJ or Imaris software.

### Quantitative analyses of microscopic images

Quantitative analyses used ImageJ and followed previously published standard methods (Patel et al., 2022; Du et al., 2018a). The fluorescence intensities for Ths:sfGFP^endo^ (Fig. 3E,E’) and Pnt:GFP (Figure 7E,E’) were measured from a selected region of interest (ROI) from maximum-intensity projections (YZ projections) encompassing the wing disc sections of fixed thickness. For comparing the intensities of Pyr:mCherry^endo^ and Ths:sfGFP^endo^ in the AMP niche (Fig.3C), the intensity values were obtained using XY sections; the respective fluorophore intensities were normalized relative to the area of the ROI. For measuring Ths:sfGFP^endo^ intensity in control and *dia-i* mutant clones (Fig.6C), GFP intensity within an ROI encompassing the clone was measured and was background corrected. Manual counting was used following (Patel et al., 2022; Du et al., 2018a) for endosomes, cytonemes, and clones from comparable 3D volume and tissue regions.

### Statistics

Statistical analyses were performed using VassarStat and GraphPad Prism 8. P-values were determined using the unpaired two-tailed t-test for pair-wise comparisons. Differences were considered significant when p-values were <0.01. All measurements were obtained from at least three independent experiments. All graphs were generated using GraphPad Prism 8 or Microsoft Excel.

## Acknowledgments

We thank Drs. T.B. Kornberg, G. M. Rubin, K. A. Guss, and S. Roth for sharing reagents; the Bloomington Stock Center and VDRC for *Drosophila* lines. This work was funded by NIH R35MG156377, R35GM124878, and R35GM124878-03S1 grants to SR.

## Author Contributions

S.R. supervised the work, designed the project, and wrote the paper. A.P. planned and performed all experiments and wrote the paper. H.H., and M.M. contributed to the genome-editing experiments and writing.

---

**Supplementary Figure 1. Characterization of Pyr and Ths chimeras. (A-B’)** Anti-dpERK-stained wing disc pouches expressing the indicated Pyr or Ths chimeras. Red and dashed arrows: MAPK pathway activation (anti-dpERK; nuclear localization) in niche-adhering AMPs that received fluorescently tagged ligands expressed from the *dpp*-zone within the wing disc pouch. **(C–D’)** AMP colonization of wing disc pouches expressing Pyr:mCherry2 under *dpp*-*Gal4*. XY and YZ (C,C’) or XY and XZ (D,D’) views as indicated. Arrowhead: Pyr:mCherry puncta in AMPs (green). Blue: phalloidin. **(E,F)** Quantification of eggs laid (E) and ovary size (F) in 7-day-old females overexpressing Ths:GFP1 or Ths:GFP2 under *ths-Gal4*, compared to *ths*-*Gal4*/+ controls (see Methods). *P*-values from two-tailed *t*-tests (relative to control). **(G,H)** Predicted structures of Thisbe (G) and Pyramus (H); color codes as indicated. Genotypes are indicated in each panel. Scale bar: 10 μm.

**Supplementary Figure 2. Generation of genome-edited *Drosophila* lines carrying *pyr:mCherry^endo^*, *ths:sfGFP^endo^*, and *htl:mCherry^endo^* knock-in constructs. (A-C)** CRISPR/Cas9-based genome editing strategy for generating *pyr:mCherry^endo^* (A), *ths:sfGFP^endo^* (B) and *htl:mCherry^endo^* (C) alleles. **(D)** Screening results for each targeted allele (also see Materials and Methods). Sequence-verified lines used in this study are indicated in row #8. See source data in FigS2D for the null rescue analyses.

**Supplementary Figure 3. Characterization of endosomal localization of Pyr:mCherry^endo^ and Ths:sfGFP^endo^ in recipient AMPs. (A)** A table showing quantitative analyses of internalized Pyr:mCherry^endo^ and Ths:sfGFP^endo^ puncta from *htl:mCherry*/+, *ths:sfGFP^endo^*/+ wing discs for their localization in early, late, and recycling endosomes of AMPs (also see Figure 4).

**Supplementary Figure 4. Exclusive presence of externalized Pyr:mCherry^endo^ and Ths:sfGFP^endo^ along niche-adhering AMP cytonemes. (A-A”’)** Comparison of the level of α-GFP^NP^-probed Ths:GFP^endo^ received on the niche-distal AMP (A,A’) and niche-proximal AMP (A”,A”’) surfaces (genotype: *ths:sfGFP^endo^/+; htl-Gal4, UAS-Lifeact:RFP/+*); *: lack of α-GFP^NP^; arrow: α-GFP^NP^. **(B)** Quantitative analyses showing % of niche-proximal AMP cytonemes localizing Pyr:mCherry^endo^ (α-Cherry^NP^) and Ths:sfGFP^endo^ (α-GFP^NP^) puncta along their surfaces (genotypes indicated). **(C-D’)** Comparison of α-GFP^NP^-probed Ths:GFP^endo^ localization in control AMP-wing disc junction (C,C’; arrow; genotype: *ths:sfGFP^endo^/+*) and *htl-*deficient AMP-wing disc junctions (D,D’; *ths:sfGFP^endo^/UAS-htl-RNAi; htl-Gal4, UAS-Lifeact:RFP/+*); *: loss of α-GFP^NP^-probed Ths:GFP^endo^ localization. **(E)** Loss (*) of α-mCherry^NP^ staining in the *pyr-*niche from *pyr:mCherry^endo^*wing discs upon the loss of niche occupancy of CD8:GFP-marked *htl-*deficient AMPs (and AMP cytonemes); genotype: *pyr:mCherry^endo^/UAS-htl-RNAi*; *htl-Gal4, UAS-CD8:GFP/+*. Scale bars: 10 µm.

**Supplementary Figure 5. Comparison of control and *pnt-i* clones. (A-C)** Comparative analyses of the number and niche-specific orientation of cytonemes from WT and *pnt*-RNAi-expressing AMP clones. Genotypes (A) and AMP position relative to the niche (A,B; see Figure 6A,F-I; 7F-F”) are indicated. **(C)** Comparative analyses of nuclear dpERK staining (pMAPK signaling activation) in WT and *pnt*-RNAi-expressing AMP clones. Genotypes and AMP position relative to the niche are indicated. (**A,C**) *: p<0.0001 (two-tailed t-test comparing control and mutant).

