## Supplementary Figures for "Cytoneme feedback ensures signaling specificity when multiple ligands converge on a common receptor"

**Supplementary Figure 1**

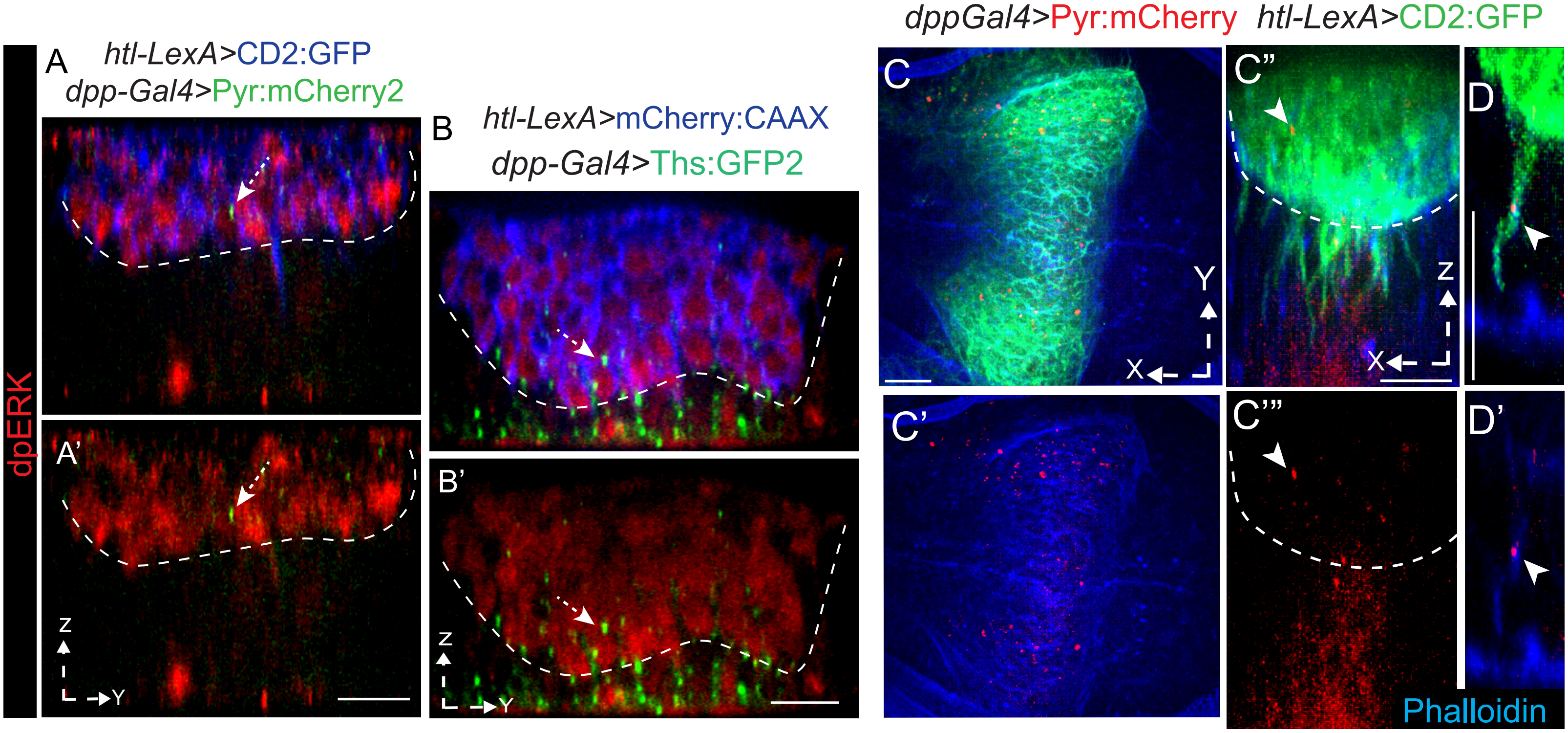


E

| Genotype | # of female in crosses | # of eggs deposited in 24 hours (No. ± S.D.) |
|---|---|---|
| *ths-Gal4/+* | 10 | 28.7 ± 4.8 |
| *ths-Gal4/UAS-*Ths:GFP1 | 10 | 27.7 ± 3.9 (p= 0.616) |
| *ths-Gal4/UAS-*Ths:GFP2 | 10 | 0.0 ± 0.0 (p<0.0001) |

F

| Genotype | #females analyzed | Average ovary size ± SD (microns) |
|---|---|---|
| *ths-Gal4/+* | 3 | 1018.0 ± 25.1 |
| *ths-Gal4/UAS-Ths:GFP1* | 3 | 1024.0 ± 24.8 |
| *ths-Gal4/UAS-Ths:GFP2* | 3 | 1402.2 ± 53.0 (p<0.0001) |

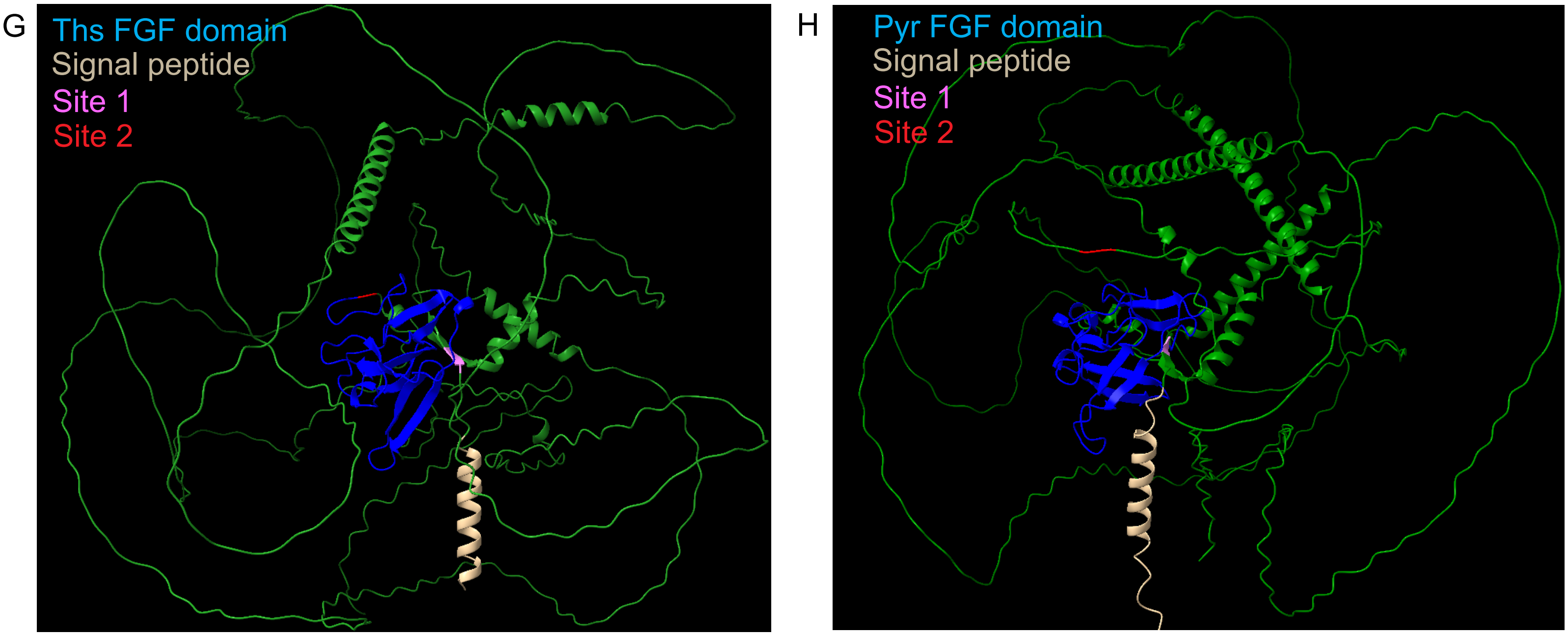

**Supplementary Figure 1. Characterization of Pyr and Ths chimeras.**

**(A-B')** Anti-dpERK-stained wing disc pouches expressing the indicated Pyr or Ths chimeras. Red and dashed arrows: MAPK pathway activation (anti-dpERK; nuclear localization) in niche-adhering AMPs that received fluorescently tagged ligands expressed from the *dpp*-zone within the wing disc pouch. **(C–D')** AMP colonization of wing disc pouches expressing Pyr:mCherry2 under *dpp-Gal4*. XY and YZ (C,C') or XY and XZ (D,D') views as indicated. Arrowhead: Pyr:mCherry puncta in AMPs (green). Blue: phalloidin. **(E,F)** Quantification of eggs laid (E) and ovary size (F) in 7-day-old females overexpressing Ths:GFP1 or Ths:GFP2 under *ths-Gal4*, compared to *ths-Gal4*/+ controls (see Methods). *P*-values from two-tailed *t*-tests (relative to control). **(G,H)** Predicted structures of Thisbe (G) and Pyramus (H); color codes as indicated. Genotypes are indicated in each panel. Scale bar: 10 μm.

### Supplementary Figure 2

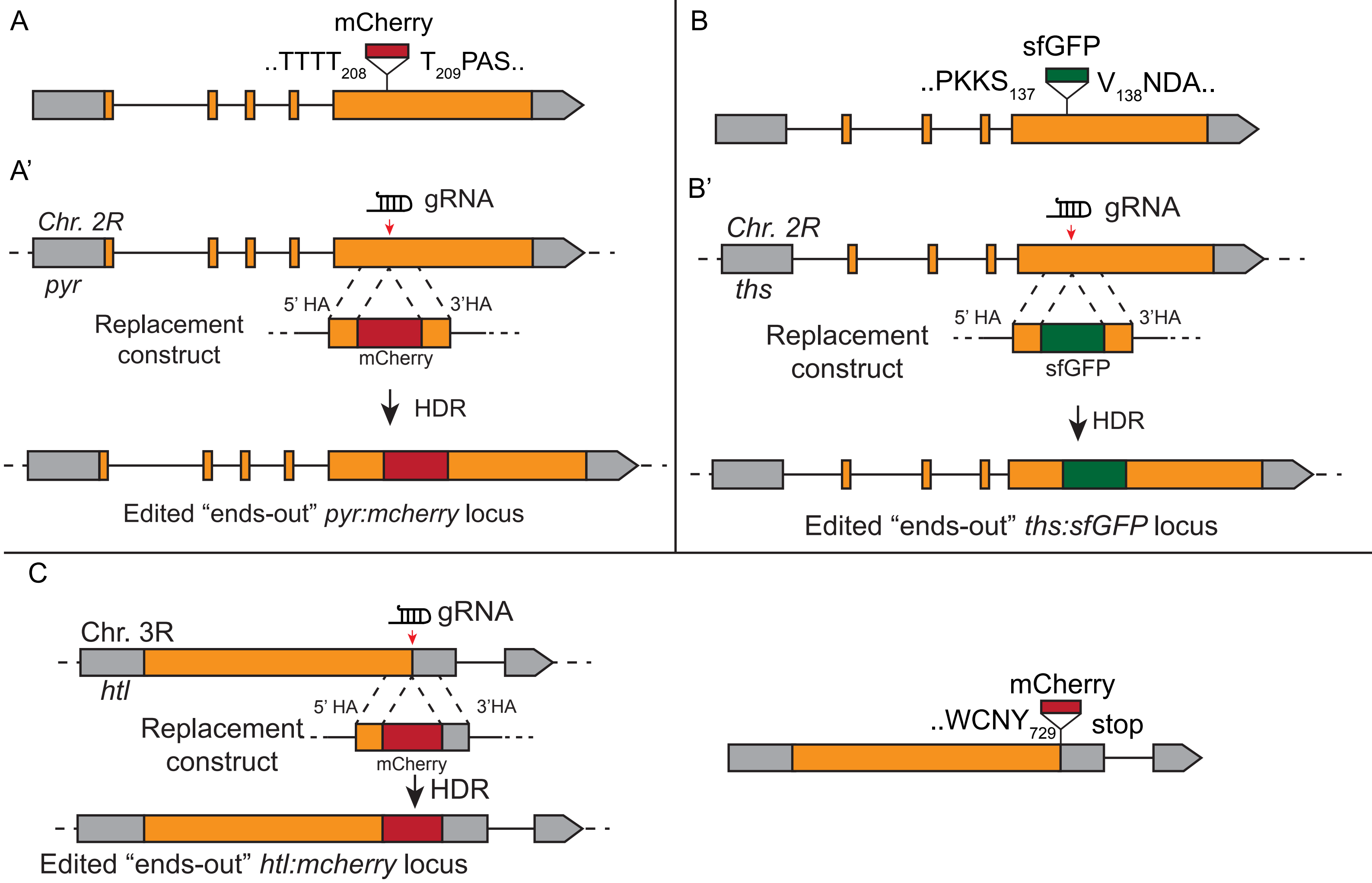


| D | *pyr:mCherry*$^{endo}$ | *ths:sfGFP*$^{endo}$ | *htl:mCherry*$^{endo}$ |
|---|---|---|---|
| (1) total fertile G0 flies | 6 | 18 | 10 |
| (2) #F1 progeny screened | 60 | 180 | 95 |
| (3) #HDR positive lines | 27 | 50 | 11 |
| (4) #ends-in HDR lines | 8 | 32 | 0 |
| (5) #ends-out HDR lines | 19 | 18 | 10 |
| (6) total normal expression and morphology from (5) | 19 | 18 | 10 |
| (7) Complete sequence verified lines from (6) | 2/19 | 3/18 | 2/10 |
| (8) #verified lines homozygous viable from (7) (line numbers used) | 2/2 (M1-10, M1-14) | 3/3 (M9-2, M18-6, F9-2) | 2/2 (F2-10, F4-8) |
| (9) Null mutants rescued by all sequenced lines from (8) (null mutant genotype) | Yes (*pyr-Gal4/CyO, DfBSC25/CyO*) | Yes (*ths-Gal4/CyO, DfBSC25/CyO, ths*$^{759}$*/CyO*) | N/A |

**Supplementary Figure 2. Generation of genome-edited *Drosophila* lines carrying *pyr:mCherry*$^{endo}$, *ths:sfGFP*$^{endo}$, and *htl:mCherry*$^{endo}$ knock-in constructs. (A-C)** CRISPR/Cas9-based genome editing strategy for generating *pyr:mCherry*$^{endo}$ (A), *ths:sfGFP*$^{endo}$ (B) and *htl:mCherry*$^{endo}$ (C) alleles. **(D)** Screening results for each targeted allele (also see Materials and Methods). Sequence-verified lines used in this study are indicated in row #8. See source data in FigS2D for the null rescue analyses.

**Supplementary Figure 3**

A

| Genotype | Average % (±S.D.) FGF-FGFR puncta in endosomal compartments | | | |
|---|---|---|---|---|
| | #Discs | α-Rab5 (early) | α-Rab7 (late) | α-Rab11 (recycling) |
| *pyr:mCherry*$^{endo}$*/+; hlt-fTRG/+* | 3 | 68.9 ± 7.7 (77 puncta) | 86.2 ± 2.4 (106 puncta) | 5.7 ± 2.5 (91 puncta) |
| *ths:sfGFP*$^{endo}$*/+; htl:mCherry*$^{endo}$ | 3 | 85.3 ± 2.2 (168 puncta) | 80.4 ± 4.9 (224 puncta) | 1.6 ± 0.2 (190 puncta) |

**Supplementary Figure 3. Characterization of endosomal localization of Pyr:mCherry$^{endo}$ and Ths:sfGFP$^{endo}$ in recipient AMPs.**

**(A)** A table showing quantitative analyses of internalized Pyr:mCherry$^{endo}$ and Ths:sfGFP$^{endo}$ puncta from *htl:mCherry*/+, *ths:sfGFP$^{endo}$*/+ wing discs for their localization in early, late, and recycling endosomes of AMPs (also see Figure 4).

**Supplementary Figure 4**

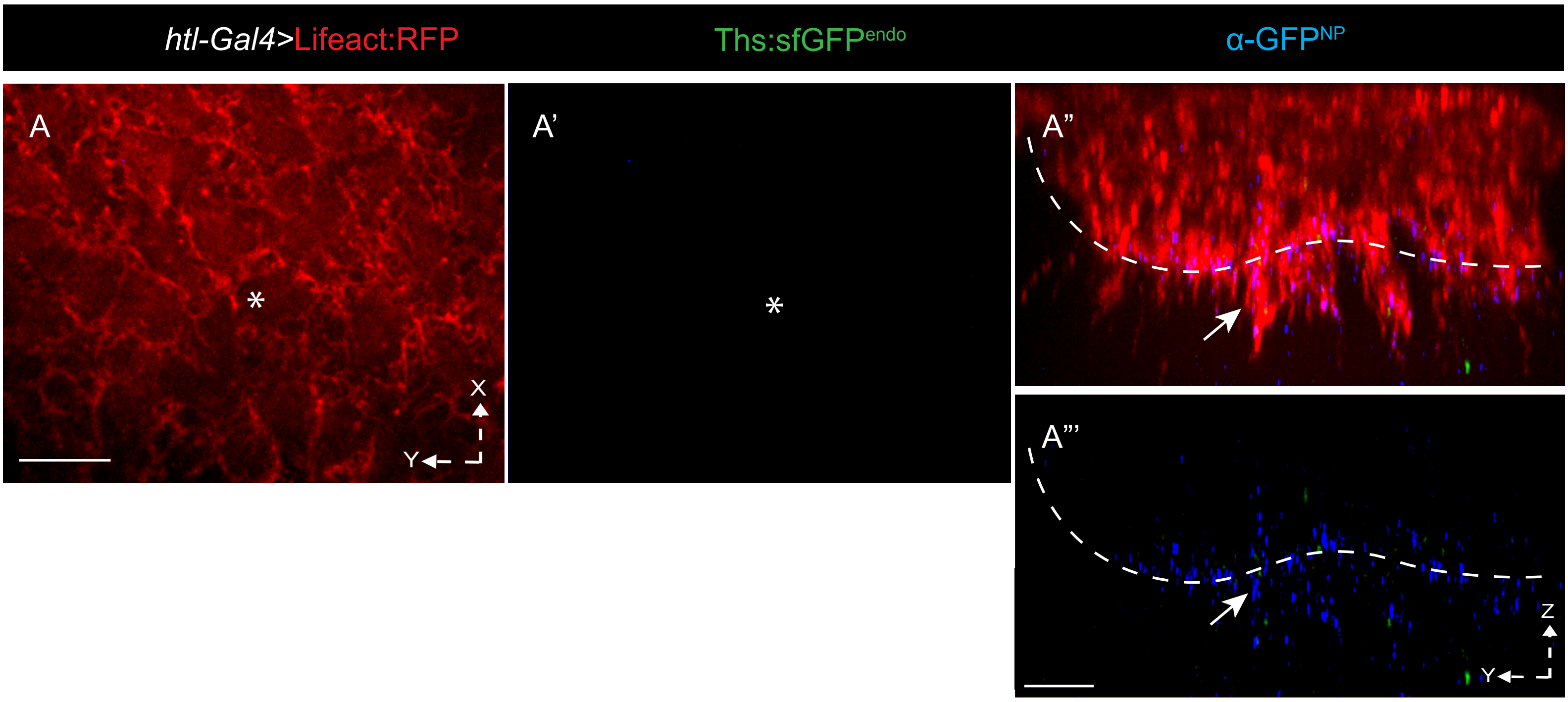


| B Genotype | #Discs | #FGFR containing cytonemes | #FGFR containing cytonemes with NP-stain | % FGFR containing cytonemes with NP-stain (±S.D.) |
|---|---|---|---|---|
| *pyr:mCherry$^{endo}$/+; hlt-fTRG/+* | 6 | 99 | 99 (α-mCherry) | 100.0 ± 0.0 |
| *ths:sfGFP$^{endo}$/+; htl:mCherry$^{endo}$* | 6 | 104 | 95 (α-GFP) | 92.5 ± 6.8 |

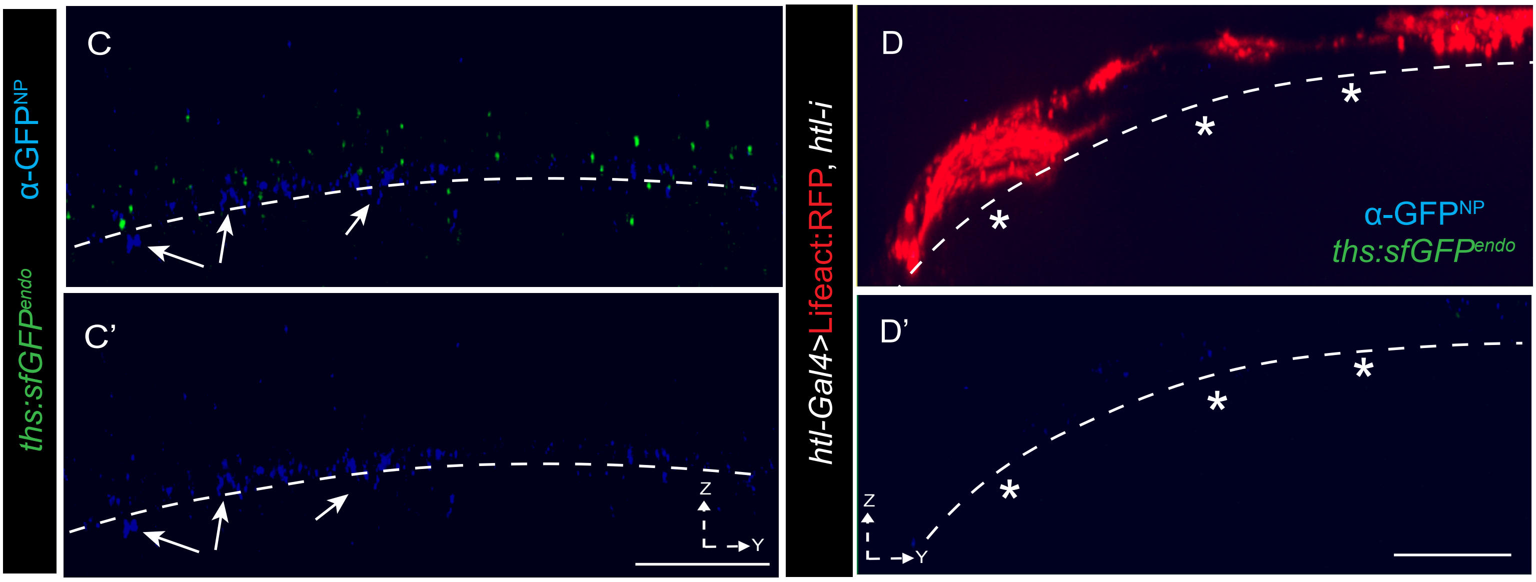


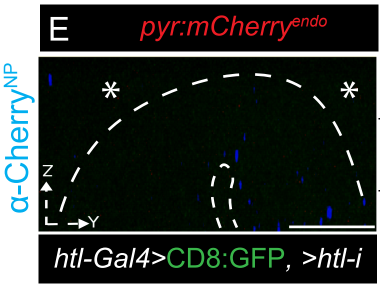

**Supplementary Figure 4. Exclusive presence of externalized Pyr:mCherry$^{endo}$ and Ths:sfGFP$^{endo}$ along niche-adhering AMP cytonemes. (A-A''')** Comparison of the level of α-GFP$^{NP}$-probed Ths:GFP$^{endo}$ received on the niche-distal AMP (A,A') and niche-proximal AMP (A'',A''') surfaces (genotype: *ths:sfGFP$^{endo}$/+; htl-Gal4, UAS-Lifeact:RFP/+*); *: lack of α-GFP$^{NP}$; arrow: α-GFP$^{NP}$. **(B)** Quantitative analyses showing % of niche-proximal AMP cytonemes localizing Pyr:mCherry$^{endo}$ (α-Cherry$^{NP}$) and Ths:sfGFP$^{endo}$ (α-GFP$^{NP}$) puncta along their surfaces (genotypes indicated). **(C-D')** Comparison of α-GFP$^{NP}$-probed Ths:GFP$^{endo}$ localization in control AMP-wing disc junction (C,C'; arrow; genotype: *ths:sfGFP$^{endo}$/+*) and *htl*-deficient AMP-wing disc junctions (D,D'; *ths:sfGFP$^{endo}$/UAS-htl-RNAi; htl-Gal4, UAS-Lifeact:RFP/+*); *: loss of α-GFP$^{NP}$-probed Ths:GFP$^{endo}$ localization. **(E)** Loss (*) of α-mCherry$^{NP}$ staining in the *pyr*-niche from *pyr:mCherry$^{endo}$* wing discs upon the loss of niche occupancy of CD8:GFP-marked *htl*-deficient AMPs (and AMP cytonemes); genotype: *pyr:mCherry$^{endo}$/UAS-htl-RNAi*; *htl-Gal4, UAS-CD8:GFP/+*. Scale bars: 10 µm.

**Supplementary Figure 5**

A

| | #discs | Clonal cell layer | Total #clones | Average disc directed cytoneme/cell | Average lateral cytoneme/cell | Average cell angle |
|---|---|---|---|---|---|---|
| *htl>FRT> Lifeact:GFP/+* (control) | 3 | Proximal | 31 | 2.2 ± 0.2 | 0.00 ± 0.00 | 85.8 ± 3.6 |
| | | Intermediate | 31 | 1.0 ± 0.2 | 1.0 ± 0.1 | 66.4 ± 7.9 |
| | | Distal | 34 | 0.00 ± 0.00 | 4.88 ± 0.5 | 0.4 ± 0.6 |
| *htl>FRT> Lifeact:GFP/ pnt-i* | 7 | Proximal | 0 * | 0.00 ± 0.00 * | 0.00 ±0.00 * | 0.00 ± 0.00 * |
| | | Intermediate | 0 * | 0.00 ± 0.00 * | 0.00 ± 0.00 * | 0.00 ± 0.00 * |
| | | Distal | 52 | 0.00 ± 0.00 | 5.5 ± 1.2 | 1.1 ± 0.8 |

B

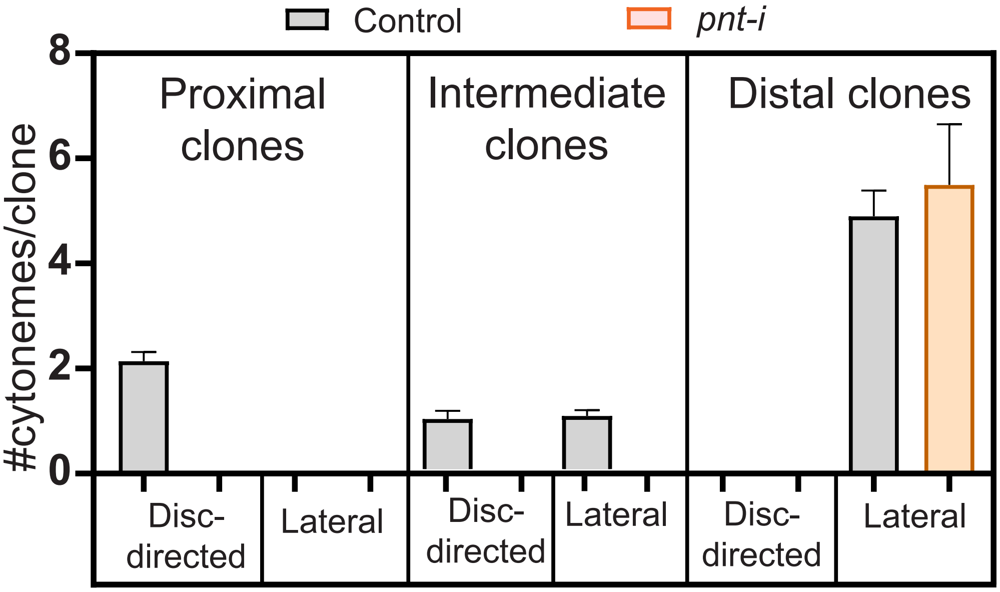


C

| | #discs | AMP layer | Total #clones | Average % clones with dpERK |
|---|---|---|---|---|
| *htl>FRT> CD8:RFP/+* (control) | 3 | Proximal | 40 | 90.2 ±2.0 |
| | | Intermediate | 45 | 51.0 ± 4.9 |
| | | Distal | 52 | 5.9 ± 0.4 |
| *htl>FRT> CD8:RFP/ pnt-i* | 4 | Proximal | 0 | 0.0 ± 0.0 * |
| | | Intermediate | 0 | 0.0 ± 0.0 * |
| | | Distal | 55 | 0.0 ± 0.0 * |

**Supplementary Figure 5. Comparison of control and *pnt-i* clones. (A-C)** Comparative analyses of the number and niche-specific orientation of cytonemes from WT and *pnt*-RNAi-expressing AMP clones. Genotypes (A) and AMP position relative to the niche (A,B; see Figure 6A,F-I; 7F-F") are indicated. **(C)** Comparative analyses of nuclear dpERK staining (pMAPK signaling activation) in WT and *pnt*-RNAi-expressing AMP clones. Genotypes and AMP position relative to the niche are indicated. (**A,C**) *: p<0.0001 (two-tailed t-test comparing control and mutant).
