## Supplementary Table 1 for "Cytoneme feedback ensures signaling specificity when multiple ligands converge on a common receptor"

### Supplementary Table 1: Reagent list used in this study

| REAGENT or RESOURCE | SOURCE/METHOD | IDENTIFIER |
|---|---|---|
| Antibodies | | |
| rabbit anti-dpERK (Phospho-p44/42 MAPK (Erk1/2) (Thr202/Tyr204) (1:250) | Cell Signaling | AB_11207064 |
| Rabbit anti-Twist (1:2000) | (Roth et al., 1989) | N/A |
| Rabbit anti-Rab7 (1:3000) | (Tanaka and Nakamura, 2008) | N/A |
| Rabbit anti-Rab11 (1:8000) | (Tanaka and Nakamura, 2008) | N/A |
| Rabbit anti-mCherry (1:1000) | Abcam | Cat# AB167453 RRID: AB_167453 |
| Rabbit anti-GFP (1:3000) | Abcam | Cat# AB6556 RRID: AB_305564 |
| Goat anti-Mouse IgG (H+L), Alexa Fluor 555 | Thermo Fisher Scientific | Cat# A21434; RRID: AB_2535855 |
| Goat anti-Mouse IgG (H+L), Alexa Fluor 647 | Thermo Fisher Scientific | Cat# A28181; RRID: AB_2536165 |
| Goat anti-Rabbit IgG (H+L), Alexa Fluor 555 | Thermo Fisher Scientific | Cat# A21428; RRID: AB_2535849 |
| Goat anti-Rabbit IgG (H+L), Alexa Fluor 647 | Thermo Fisher Scientific | Cat# A21244; RRID: AB_2535812 |
| Bacterial and Virus Strains | | |
| DH5 Alpha | | |
| Chemicals, Peptides, and Recombinant Proteins | | |
| Phalloidin iFlor 555 | Abcam | Cat# ab176759 |
| Phalloidin iFlor 647 | Abcam | Cat# ab176756 |
| Critical Commercial Assays | | |
| CloneJET PCR Cloning Kit | Thermo Fisher Scientific | Cat# K1231 |
| Zymoclean Gel DNA Recovery Kit | Zymo Research | Cat# D4007 |
| GeneJET Plasmid Miniprep Kit | ThermoFisher Scientific | Cat# K0502 |
| GeneJET Plasmid Midiprep Kit | ThermoFisher Scientific | Cat #K0481 |
| 2X PCR Premix | Syd Labs | Cat# MB067-EQ2R-L |
| Deposited Data | | |
| Raw data from all the figures | This paper | |
| Experimental Models: Organisms/Strains | | |

| | | |
|---|---|---|
| *D. melanogaster*: *UAS-CD8:GFP* | BDSC | BDSC_5130 |
| *D. melanogaster*: *UAS-CD8:RFP* | BDSC | BDSC_32218 |
| *D. melanogaster*: *UAS-Lifeact:RFP* | BDSC | BDSC_58362 |
| *D. melanogaster*: *UAS-mCherryCAAX* | BDSC | BDSC_59021 |
| *D. melanogaster*: *lexO-mCherryCAAX* | (Roy et al., 2014) | N/A |
| *D. melanogaster*: *lexO-CD2:GFP* | BDSC | BDSC_66544 |
| *D. melanogaster*: *UAS-Lifeact:GFP* | BDSC | BDSC_57326 |
| *D. melanogaster*: *UAS-diaRNAi* | BDSC | BDSC_33424 |
| *D. melanogaster*: *htl-Gal4* | BDSC | BDSC_40669 |
| *D. melanogaster: pnt:GFP* | BDSC | BDSC_42680 |
| *D. melanogaster*: *UAS-pntRNAi* | BDSC | BDSC_35038 |
| *D. melanogaster*: *UAS-pntP1* | BDCS | BDSC_869 |
| *D. melanogaster: ths-Gal4* | BDSC | BDSC_77475 |
| *D. melanogaster*: *{nos-Cas9}ZH-2A* | BDSC | BDSC_54591 |
| *D. melanogaster*: *hs-Flp* | BDSC | BDSC_6 |
| *D. melanogaster*: $w^{1118}$ | BDSC | BDSC_3605 |
| *D. melanogaster*: $htl{:}GFP^{fTRG}$ | VDRC | 318120; Flybase_FBst0491541 |
| *D. melanogaster*: *UAS-htlRNAi* | VDRC | 6692; Flybase_FBst0470407 |
| *D. melanogaster: dpp-Gal4* | (Roy et al., 2014) | N/A |
| *D. melanogaster*: *htl-LexA* | (Patel et al., 2022) | N/A |
| *D. melanogaster*: *pyr-Gal4* | (Patel et al., 2022) | N/A |
| *D. melanogaster*: *htl>FRT>stop>FRT>Gal4* | (Patel et al., 2022) | N/A |
| *D. melanogaster*: *UAS-Pyr:GFP1* | This study | N/A |
| *D. melanogaster*: *UAS-Pyr:sfGFP2* | (Patel et al., 2022) | N/A |
| *D. melanogaster: UAS-Ths:sfGFP1* | This study | N/A |
| *D. melanogaster*: *UAS-Ths:sfGFP2* | (Patel et al., 2022) | N/A |
| *D. melanogaster*: *UAS-Pyr:mCherry2* | This study | N/A |
| *D. melanogaster*: $pyr{:}mCherry^{endo}$ | This study | N/A |
| *D. melanogaster*: $ths{:}sfGFP^{endo}$ | This study | N/A |
| *D. melanogaster*: $htl{:}mCherry^{endo}$ | This study | N/A |
| Oligonucleotides | | |
| ***pJet-pyr:mCherry*** **(**$pyr{:}mCherry^{endo}$**)** | | |
| GTCGTGACCGCCACTGGGCTAGC | This study | gRNA F P3 pCFD3 |
| AAACGCTAGCCCAGTGGCGGTCA | This study | gRNA R P3 pCFD3 |
| ACACCAGCATCCCCAGTGGCGGTCACAAAAC | This study | C_F_P3 with gRNA change |
| CACTGGGGATGCTGGTGTCTTGTACAGCTCGTCCATGCC | This study | Cherry_Covrhng_R_P3_with gRNA change |
| CACAACAACAACAACCACAACAACCATGGTGAGCAAGGGCGAG | This study | Cherry_Novrhng_F_P3 |
| CGAACTCGTGGCCGTTCAC | This study | cherry_Nter_seq_R |

| | | |
|---|---|---|
| CCCTGAAGGGCGAGATCAAG | This study | cherrymidseqF |
| GACCTCAGCGTCGTAGTGGC | This study | cherrymidseqR |
| GGTTGTTGTGGTTGTTGTTGTTGTG | This study | N_Cter_P2_R |
| CCCAGTCACGACGTTGTAAAACG | This study | Primer 1 P3endo |
| AATTCGAGCTCGGTACCGGGTTAAACCCTTTC GCCCAG | This study | Primer 2 P3endo |
| GGTACCGAGCTCGAATTCACTGG | This study | Primer 3 P3endo |
| CATAGAAACCTCTAGAACTTTTAAAGTTTTGC | This study | Primer 4 P3endo |
| AGAAACGCCACAAAGGAGTGC | This study | Primer 5 P3endo |
| GCACTCCTTTGTGGCGTTTCTCTATAAAATAAA AAAAAAGACAAAGGGAAAGAGAACTTAGTAAG | This study | Primer 6 P3endo |
| GCGATTTCTCTGCAGGATCACC | This study | Primer 7 P3endo |
| TGCAGTTGCAGCTGCCG | This study | Primer 8 P3endo |
| GCGTTAGTGTTAATCATGCAATACCTGC | This study | primer 9 P3endo |
| CACACAGGAAACAGCTATGACC | This study | Primer 10 P3endo |
| TCGTTCGGATAGCTATCGTCGATTATATTATTAG CATCAATATGATTGCGATTCTGC | This study | pyr614-633R |
| GCCCAAGGCCACCTCCTC | This study | PyrMidSeqF |
| GAGGAGGTGGCCTTGGGC | This study | PyrMidSeqR |
| ***pJet-ths:sfGFP (ths:sfGFP$^{endo}$)*** | | |
| GTCGCGTGTCGGGTTCACGCACCG | This study | gRNA F T2 pCFD3 |
| AAACCGGTGCGTGAACCCGACACG | This study | gRNA R T2 pCFD3 |
| GCCAAGCTTGCATGCCCCGTTAGTGTGTTGTT CAGGTTGC | This study | C_pUC19ovrhng _R_T2 |
| CAACATCTGACGACAGTATTAGCAGC | This study | Ends in_out_F_T2 |
| CCATATTACGGCGACCTGGC | This study | genome F T2 |
| AATTCGAGCTCGGTACCCCATATTACGGCGAC CTGGC | This study | genome_pUC19 ovrhng_F_T2 |
| CAGACGGCAAACTAAACGTTCACTC | This study | gRNAseqF1 ThsCRISPR |
| CCACCATTGTGAATGCCAGGG | This study | gRNAseqF2 ThsCRISPR |
| GTGATGATGTCGCACCTGATGTTG | This study | gRNAseqR1 ThsCRISPR |
| GGACTAGCTGATACTCGTTGTTGGTG | This study | gRNAseqR2 ThsCRISPR |
| CGACTCACTATAGGGAGAGCGGC | This study | pJET1.2 forward sequencing primer |
| AAGAACATCGATTTTCCATGGCAG | This study | pJET1.2 reverse sequencing primer |
| GTTGATGTTCTTGTCCACGCTGAAG | This study | Rev2 R T2 |
| CTGCTATCAGCGGACCGATC | This study | Rev3 R T2 |

| | | |
|---|---|---|
| GTCGAGGGGCAAGGCGGCATGTCCAAGGGC GAGGAGC | This study | sfGFP_linker_F_ T3 |
| ATGTCCAAGGGCGAGGAGC | This study | sfGFP_Nter_F |
| TCACAGAGGTAAGCCTGTGGGGC | This study | gRNAmut F |
| CACAGGCTTACCTCTGTGAGTGAACCCGACA CGTCGC | This study | gRNAmut R |
| GAAGCAGCACGATTTCTTCAAGAGCG | This study | sfgfpmidF check |
| CGCTCTTGAAGAAATCGTGCTGCTTC | This study | sfgfpmidR check |
| GCTGCGACGACAGCAACAC | This study | Ths MidSeqF |
| GTGTTGCTGTCGTCGCAGC | This study | Ths MidSeqR |
| ***pUC19-htl:mCherry$^{endo}$ (htl:mCherry$^{endo}$)*** | | |
| GTCGTGTAATTATTAAACGAATC | This study | gRNA Htl Cherry F |
| AAACGATTCGTTTAATAATTACAC | This study | gRNA Htl Cherry R |
| GTACAAGTAAACGAATCAGGATCCTTAGATGA GATCAG | This study | C_F_HtlCherry |
| GCCAAGCTTGCATGCCCCAAATGCCAACAGCT AAACTCCAG | This study | C_puc19ovrhng_ R_HtlCherry |
| GGATCCTGATTCGTTTACTTGTACAGCTCGTC CATGCC | This study | cherry_Covrhng_ R_HtlCherry |
| CTTCCCCGAGGGCTTCAAGTG | This study | cherry midseqF |
| CACTTGAAGCCCTCGGGGAAG | This study | cherry midseqR |
| CGCCTTTGGTGCAATTTTGG | This study | gRNA seqR HtlCherry |
| GGATCTGATCAAATTTGCCCACC | This study | Htl_midseq_F |
| CTGACGAGCACATTCCTGGC | This study | Htl_midseq_R |
| TGTAAAACGACGGCCAGT | This study | M13F |
| CAGGAAACAGCTATGAC | This study | M13R |
| GAGGCTCATTGCGAATAAATCGGAC | This study | rev2 HtlCherry |
| AATTCGAGCTCGGTACCTGGATCTTTGTGCCC TGCCATG | This study | N_pUC19ovrhng _F_HtlCherry |
| ***UAS-Ths:sfGFP1*** | | |
| GCCAAGCTTGCATGCCTCTAGACTACGCAAAT CTCTGATGAGTGAACC | This study | C- puc19overhng_R _T1 |
| CATGGATGAACTATACAAAGTAATAATGTCACA GTGTGCACACAACAAAG | This study | C_GFPovrhng_F _T1 |
| GCACACTGTGACATTATTACTTTGTATAGTTCAT CCATGCCATGTGTAATC | This study | GFP_Covrhng_R _T1 |
| TATGTACAGTAGAAGATTACATGAGTAAAGGAG AAGAACTTTTCACTGG | This study | GFP_Novrhng_F _T1 |
| GCGATGGCCCTGTCCTTTTAC | This study | GFPseqF |
| GTAAAAGGACAGGGCCATCGC | This study | GFPseqR |
| AATTCGAGCTCGGTACCATGTCGAATCAGTTA GAGAGACTGCTG | This study | N- pUC19ovrhng_F _T1 |
| AGTTCTTCTCCTTTACTCATGTAATCTTCTACTG TACATAATGCACCTGAAATC | This study | N_GPFovrhng_R _T1 |

| | | |
|---|---|---|
| ***UAS-Pyr:GFP1*** | | |
| GCATGGATGAACTATACAAAACATTGAGCAAACACAGCGAAC | This study | C_GFPovrhng_F_P1 |
| GCCAAGCTTGCATGCCGGATCCCTCGAGCTATAAATCTATATAATACAAGCTAACAAAATACTTACCAC | This study | C_pUC19ovrhng_R_P1 |
| TCGCTGTGTTTGCTCAATGTTTTGTATAGTTCATCCATGCCATGTGTAATCC | This study | GFP_Covrhng_R_P1 |
| GCGCCGCGAAAAATGTTTTAATGAGTAAAGGAGAAGAACTTTTCACTGG | This study | GFP_Novrhng_F_P1 |
| AGTTCTTCTCCTTTACTCATTAAAACATTTTTCGCGGCGCTTG | This study | N_GFPovrhng_R_P1 |
| AATTCGAGCTCGGTACCGCGGCCGCATGTTCCACAAGTTCATGCCCAATG | This study | N_pUC19ovrhng_F_P1 |
| ***UAS-Pyr:mCherry2*** | | |
| GGTTGTTGTGGTTGTTGTTGTTGTG | This study | N_Cter_P2_R |
| CACAACAACAACAACCACAACAACCATGGTGAGCAAGGGCGAG | This study | Cherry_Novrhng_F_P3 |
| Recombinant DNA | | |
| pUC19 | Addgene | 50005 |
| pUASt | DGRC | 1000 |
| pCFD3 | (Port et al., 2014; Port et al.) | N/A |
| pCFD3-pyr:mCherry-gRNA | This study. The “gRNA F P3 pCFD3” and “gRNA R P3 pCFD3” primers were used to clone the gRNA in pCFD3 using the protocol outlined in (Du et al., 2018b) . | N/A |
| pCFD3-ths:sfGFP-gRNA | This study. The “gRNA F T2 pCFD3” and “gRNA R T2 pCFD3” primers were used to clone the gRNA in pCFD3 using the protocol outlined in (Du et al., 2018b). | N/A |
| pCFD3-htl:mCherry-gRNA | This study. The “gRNA Htl Cherry F” and “gRNA Htl Cherry R” primers were used to clone the gRNA in pCFD3 using the protocol | N/A |

| | | |
|---|---|---|
| | outlined in (Du et al., 2018b) | |
| pJet-pyr:mCherry$^{endo}$ | This study. The 5' homology arm (HA) containing 1.0kb of *pyr* genomic region upstream of the insertion site, and the 3’ HA containing 1.2kb of *pyr* genomic region downstream of the insertion site were amplified from the genomic DNA (gDNA) of the *nos-Cas9* fly and were assembled with a PCR amplified mCherry amplicon into the pJet1.2 vector using Gibson Assembly. To prevent gRNA-mediated Cas9 retargeting of the edited genome, we introduced synonymous mutations in 3’HA using “C_F_P3 with gRNA change” and “Cherry_Covrhng_R_P3_with gRNA change” primers. | N/A |
| pJet-ths:sfGFP$^{endo}$ | This study. The 5' homology arm (HA) containing 1.5kb of *ths* genomic region upstream of the insertion site, and the 3’ HA containing 1.5kb of *ths* genomic region downstream of the insertion site were amplified from the genomic DNA (gDNA) of the *nos-Cas9* fly and were assembled with a | N/A |

| | | |
|---|---|---|
| | PCR amplified "VEGQGG-sfGFP-GSGGGS" amplicon into the pJet1.2 vector using Gibson Assembly. To prevent gRNA-mediated Cas9 retargeting of the edited genome, we introduced synonymous mutations in 5'HA using "gRNAmut F" and "gRNAmut R" primers. | |
| pUC19-htl:mCherry$^{endo}$ | This study. The 5' homology arm (HA) containing 1.5kb of *htl* genomic region upstream of the insertion site, and the 3' HA containing 1.5kb of *htl* genomic region downstream of the insertion site were amplified from the genomic DNA (gDNA) of the *nos-Cas9* fly and were assembled with a PCR amplified mCherry amplicon into the pUC19 vector using Gibson Assembly. Insertion of mCherry disrupts gRNA sequence. | N/A |
| pUAST-Pyr:GFP1 | This study. An ectopic sfGFP sequence was inserted in frame between Leucine-33 and Threonine-34 (site 1) of the original 766 amino-acid-long Pyr using Gibson assembly (Figure 1B). | N/A |

| | | |
|---|---|---|
| | The 144bp N-terminal fragment form the start codon to the insertion site was amplified using “N_pUC19ovrhng_F_P1” and “N_GFPovrhng_R_P1” primers; the 754bp GFP fragment was amplified using “GFP_Novrhng_F_P1” and “GFP_Covrhng_R_P1” primers; the 2250bp C-terminal fragment was amplified using “C_GFPovrhng_F_P1” and “C_pUC19ovrhng_R_P1” primers. The three fragments were assembled in pUC19 vector using Gibson assembly and cloned into *pUAST* using Not1/Xho1 restriction sites. | |
| pUAST-Ths:sfGFP1 | This study. In *UAS-Ths:sfGFP1*, a sfGFP sequence was inserted in frame between Tyrosine-25 and Valine-26 (site 1) of the original 748 amino-acid-long Ths using Gibson assembly (Figure 1B).<br><br>The 112bp N-terminal fragment (112 bp) form the start codon to the insertion site was amplified using “N_pUC19ovrhng_F | N/A |

| | | |
|---|---|---|
| | _T1" and "N_GFPovrhng_R_T1" primers; the sfGFP fragment (754 bp) was amplified using "GFP_Novrhng_F_T1" and "GFP_Covrhng_R_T1" primers; the 2213bp C-terminal fragment was amplified using "C_GFPovrhng_F_T1" and "C_pUC19ovrhng_R_T1" primers. The three fragments were assembled in pUC19 vector using Gibson assembly and cloned into *pUAST* using Kpn1/Xba1 restriction sites. | |
| pUAST-Pyr:mCherry2 | This study. An ectopic mCherry sequence was inserted in frame between Threonine-208 and Threonine-209 (site 2) of the original 766 amino-acid-long Pyr using Gibson assembly (Figure 1B).<br><br>The 649bp N-terminal sequence from the start codon to Threonine-208 was amplified using "N_pUC19ovrhng_F_P2" and "N_Cter_P2_R" primers; mCherry sequence (760 bp) was amplified using "Cherry_Novrhng_F_P3" and "sfGFP_Nhe1_pUC | N/A |

| | | |
|---|---|---|
| | ovrhng_R_P2" primers. Both fragments were assembled in pUC19 using Gibson assembly and cloned into the pre-existing *pUAST-pyr:sfGFP2* ((Patel et al., 2022) using Not1/Nhe1 restriction sites. | |
| pHtl-enh-FRT-stop-FRT3-FRT-FRT3-Gal4 | (Patel et al., 2022) | N/A |
| pBP-htl-enh-nlsLexA::p65Uw | (Patel et al., 2022) | N/A |
| Software and Algorithms | | |
| Fiji- ImageJ 1.52p | ImageJ | https://fiji.sc |
| Prism 8.0 | GraphPad | https://www.graphpad.com/ |
| Adobe Photoshop 22.5.1 | Adobe | https://www.adobe.com |
| Adobe Illustrator 25.4.1 | Adobe | https://www.adobe.com |
| Microsoft Excel (Version 2111) | Microsoft | https://www.office.com |
| SnapGene 3.3.4 | SnapGene | https://www.snapgene.com |
| VassarStats | | vassarstats.net |
| Imaris 9.5.0 | Imaris | https://imaris.oxinst.com |
| Andor iQ3 | Oxford Instruments | https://andor.oxinst.com/products/iq-live-cell-imaging-software/ |
| AlphaFold Protein Structure Database | EMBL-EBI | https://alphafold.ebi.ac.uk/ (Jumper et al., 2021; Varadi et al., 2021) |
| ChimeraX 1.9 | Resource for Biocomputing, Visualization, and Informatics at the University of California, San Francisco | https://www.rbvi.ucsf.edu/chimerax/ (Meng et al., 2023) |
| Grammarly | Grammarly | https://www.grammarly.com/ |
| Zen 3 | Carl Zeiss Microscopy GmbH | https://www.zeiss.com/microscopy |

| | | |
|---|---|---|
| | | /int/home.html?vaURL=www.zeiss.com/microscopy |
